# Pathogenic mutations in ATAD3A cause dysregulation of RagC/D-TFEB axis and disrupt lysosomal homeostasis

**DOI:** 10.64898/2026.09.23.753918

**Authors:** Mathew B. McDougal, Abigail Sandoval, Mike Kinter, Antrix Jain, Sukyeong Lee, Sung Yun Jung, Wan Hee Yoon

## Abstract

We previously discovered that a *de novo* variant p.R528W in *ATAD3A*, encoding a mitochondrial membrane-anchored protein, causes a human neurological syndrome. While *ATAD3A* mutations induce aberrant lysosomal expansion accompanied by undigested material in the lysosomes, how mutant *ATAD3A* disrupts lysosomal homeostasis and whether this contributes to neurodevelopmental defects remain unknown. Here we show that pathogenic ATAD3A p.R528W expression disrupts the mTORC1-TFEB axis as revealed by dysregulation of mTORC1 substrate phosphorylation, TFEB nuclear localization, and CLEAR gene activation associated with lysosomal biogenesis. ATAD3A binds to lysosome-localized Rag C/D GTPases, which constitute a platform for TFEB recruitment, with pathogenic variants increasing this association and thereby decreasing lysosomal localization of Rag GTPases. Importantly, overexpression of *RagC* or *RagD* restores TFEB phosphorylation in human cells expressing p.R528W, and *RagC-D* overexpression or *TFEB/Mitf* knockdown rescues lysosomal expansion and neurodevelopmental defects in *Drosophila*. These data indicate that disrupted Rag GTPase recruitment to lysosomes and subsequent aberrant TFEB/Mitf activation contribute to neurodevelopmental and lysosomal phenotypes caused by pathogenic mutations in *ATAD3A*. Our work reveals a novel role for the mitochondrial resident protein ATAD3A in modulating lysosomal homeostasis through regulation of the mTORC1-TFEB axis, providing a mechanistic link between impaired mitochondrial and lysosomal homeostasis in a neurodevelopmental disorder.

## INTRODUCTION

Mitochondria and lysosomes are essential organelles for cellular function and life in metazoans. Mitochondria are centers of metabolism in the cell, while lysosomes act as a cellular hub of catabolism and nutrient sensing. Dysfunction of either organelle, including genetic mutations in genes associated with either organelle, is widely implicated in neurodevelopmental and neurodegenerative disorders.^1, 2^ While the mechanisms underlying defects arising within individual organelles have been extensively studied, emerging evidence shows that impairment of functions in one organelle often disrupts those in the other, which may accelerate disease progression.^3^ For example, loss of mitochondrial proteins such as Pink1 or Opa1 impairs lysosomal activity, whereas loss of lysosomal GBA1, a Parkinson’s disease-associated glucocerebrosidase, causes defective mitochondrial metabolism.^4, 5^ Yet, how dysfunction in one organelle propagates to disrupt homeostasis of another, and whether targeting these secondary effects can mitigate cellular stress and disease, remains largely unexplored.

The ATPase family AAA-domain-containing protein 3A (ATAD3A) is a member of the hexmeric AAA+ ATPase family, initially identified as a mitochondrial membrane-anchored protein involved in mitochondrial DNA (mtDNA) stability,^6^ mitochondrial membrane dynamics,^7^ and cholesterol metabolism.^8–11^ More recent studies have demonstrated that ATAD3A is involved in the maintenance of mitochondrial cristae,^12–16^ trafficking nucleoids,^17^ maintaining mitochondrial ribosome (mitoribome) stability,^13^ and proper folding and maintenance of electron transport chain (ETC) proteins for oxidative phosphorylation.^15, 18–20^ ATAD3A possesses a unique topology with its C-terminal AAA+ domain in the matrix and N-terminus exposed beyond the outer membrane, enabling regulation of biological processes within the mitochondria as well as cross-organelle signaling.^21^ For example, ATAD3A affects mitochondrial membrane dynamics through its N-terminal interaction with Drp1, a mitochondrial fission factor.^22^ Furthermore, the N- terminus of ATAD3A physically interacts with and inhibits protein kinase R–like endoplasmic reticulum kinase (PERK) activity to preserve local mitochondrial protein expression under ER stress conditions.^23^ These findings open the possibility that ATAD3A may play additional roles in mitochondria-organelle crosstalk.

Consistent with the key role for ATAD3A in many cellular processes, loss of function studies in mouse, nematode, fruit fly, and zebrafish models have demonstrated an essential role for *ATAD3A* in normal development.^15, 16, 24–28^ Moreover, genetic variants in *ATAD3A* have been reported to be associated with mitochondrial and neurological disorders in humans. To date, the allelic spectrum of *ATAD3A*-associated disorders comprises null, hypomorphic, and dominant-negative alleles.^9, 10, 12, 19, 26, 29^ Our group identified a recurrent de novo variant in *ATAD3A* (GenBank: NM_001170535.1; c.1582C>T: p.Arg528Trp, hereafter referred to as p.R528W) that causes a distinct neurological syndrome (Harel-Yoon syndrome/HAYOS, MIM: 617183), characterized by developmental delay, hypotonia, axonal neuropathy, and/or optic atrophy and cardiomyopathy.^12^ Additional missense variants, including p.Gly355Asp and p.Arg466Cys, inherited in an autosomal dominant manner, were subsequently identified as pathogenic mutations that cause optic atrophy, neuropathy, and cerebral palsy.^10, 29^ In addition, biallelic deletions in *ATAD3A* have been reported to result in an infantile-lethal presentation with pontocerebellar hypoplasia, hypotonia, and respiratory insufficiency syndrome [PHRINL, MIM: 618810],^9, 12^ indicating an essential role for ATAD3A in human development. *ATAD3A* is now recognized as one of the most frequently mutated genes in neonatal mitochondrial disease.^19^ However, the underlying mechanisms by which mutations in *ATAD3A* lead to defects in neurodevelopment remain incompletely understood.

For evaluating the consequences of pathogenic variants in *ATAD3A* in vivo, we have developed a robust genetic system in *Drosophila.*^10, 12, 26^ Our previous *Drosophila* functional studies demonstrated that all known heterozygous pathogenic variants, including p.R528W, act as dominant-negative mutations, disrupt mitochondrial cristae, and promote abnormal mitophagy.^12^ Interestingly, utilizing *Drosophila* models along with patient fibroblasts, we showed that the dominant *ATAD3A* pathogenic variants cause expansion of the lysosomal pool and accumulation of membranous whorls in lysosomes.^10^ Similarly, prior studies report increased lysosomal content in patient iPSC-derived neurons carrying *ATAD3A* mutations.^29^ Together, these findings indicate that dysfunction of mitochondrial protein ATAD3A may disrupt regulation of lysosomal biogenesis and function.

Lysosomal biogenesis is regulated by a conserved transcription program orchestrated by the MiT/TFE (microphthalmia/transcription factor E) family transcription factors, including transcription factor EB (TFEB) and Transcription Factor Binding To IGHM Enhancer 3 (TFE3).^30, 31^ Under nutrient-replete conditions, mTORC1-mediated phosphorylation retains TFEB and TFE3 in the cytoplasm.^32^ In contrast, nutrient deprivation or lysosomal stress inhibits mTORC1 and/or activates phosphatases that target TFEB/TFE3 for dephosphorylation, subsequent nuclear translocation, and induction of Coordinated Lysosomal Expression and Regulation (CLEAR) genes, which encode lysosomal and autophagy proteins.^31, 33–35^ Unlike canonical mTORC1 substrates such as S6K and 4E-BP, which are recruited through Raptor within the mTORC1 complex itself, TFEB and TFE3 require RagC or RagD GTPases for their recruitment to lysosomal surface and phosphorylation by mTORC1.^36, 37^ The Rag GTPases are anchored to the lysosomal surface by the Ragulator complex,^38^ and disruption of RagC/D functions or localization results in failure of mTORC1-directed phosphorylation of TFEB/TFE3, promoting nuclear localization, demonstrating RagC/D as critical regulators for TFEB/TFE3 activity.^37, 39^ Although ATAD3A dysfunction has been implicated in lysosomal biogenesis and mTORC1 signaling,^10, 29, 40^ the mechanisms by which *ATAD3A* mutations perturb lysosomal homeostasis, in particular whether *ATAD3A* pathogenic variants disrupt mTORC1 signaling, TFEB regulation, or RagC/D localization and function remain to be determined.

Here, we show the pathogenic variant ATAD3A p.R528W (*Drosophila* Atad3a/dAtad3a p.R534W), which is predicted to disrupt ATPase function, results in neurodevelopmental defects and lethality when expressed in neurons of the fruit fly. Remarkably, these phenotypes are rescued by ectopic expression of *RagC-D*. Through immunoprecipitation mass spectrometry (IP-MS) and co-IP experiments, we demonstrate that ATAD3A interacts with RagC and RagD GTPases, with ATAD3A p.R528W exhibiting a stronger interaction. Expression of ATAD3A p.R528W in human cells disrupts the mTORC1-TFEB axis with altered mTORC1 substrate phosphorylation, TFEB nuclear localization, and CLEAR gene expression. Through lysosomal IP (Lyso-IP), we discovered that ATAD3A p.R528W expression remodels the lysosomal proteome, including a reduction in Rag GTPases. Importantly, we show that ectopic expression of Rag C-D rescues Atad3a p.R534W-mediated lethality as well as lysosomal expansion phenotypes. Furthermore, knockdown of *Mitf*, the *Drosophila* homolog of TFEB, also rescues lysosomal expansion and Atad3a p.R534W-mediated lethality. Together, our data support the model that ATAD3A p.R528W disrupts the RagC/D-TFEB axis and lysosomal homeostasis, leading to the neurological pathogenesis observed in Harel-Yoon syndrome, and establishes a novel role for ATAD3A in maintaining lysosomal homeostasis.

## RESULTS

### Dominant pathogenic variants in ATAD3A predicted to disrupt ATP binding cause defects in neurodevelopment in *Drosophila*

To provide insight into the molecular mechanisms by which pathogenic ATAD3A variants impair the ATAD3A function, we generated a 3D structural model of the human ATAD3A AAA+ domain (GenBank: NP_001164006.1) based on the AlphaFold predicted ATAD3A structure to assess their potential effects on a functionally important region.^41^ The bacterial ClpB ATPase domain structure (PDB ID: 4FCV, chain A) was utilized as a solved protein structure to improve modeling. Using the program Coot, we introduced p.R528W as well as p.Gly355Asp (p.G355D), another dominant pathogenic variant associated with severe development and neurological manifestations in humans.^12, 29, 42^ AAA+ ATPases are composed of several key motifs, including the Walker A (WA) and Sensor 2 motifs that are essential for ATP binding, as well as Walker B (WB) and Sensor 1 motifs which are crucial for ATP hydrolysis (**Figure 1a-b**).^43^ Notably, the pathogenic variants p.R528W and p.G355D are located within the Sensor 2 and WA motif, respectively (**Figure 1a-b**).^43^ Indeed, the model predicts that these mutations compromise ATP binding, consistent with the finding that the p.G355D mutation reduces ATPase activity.^29^

**Figure 1.**
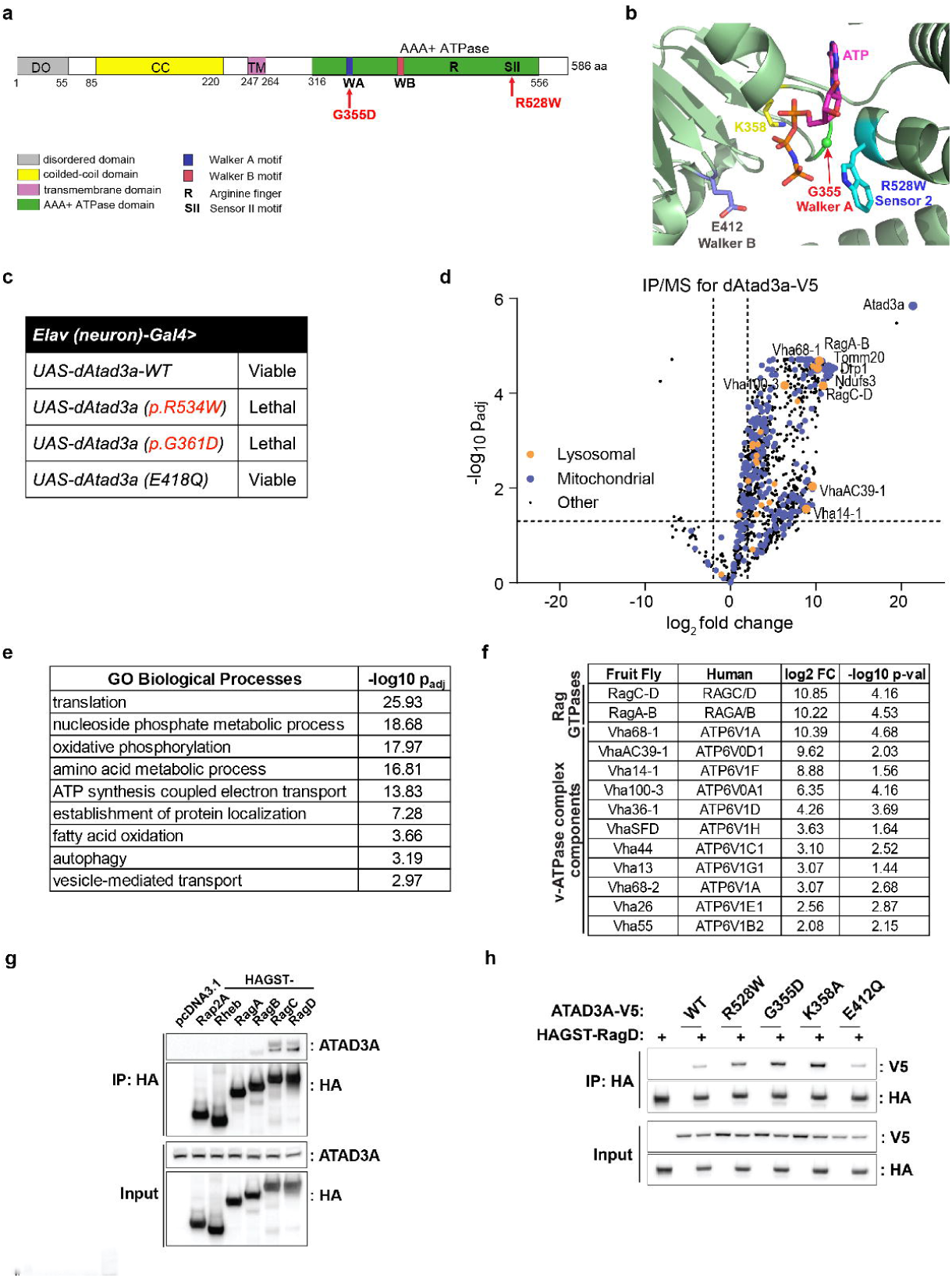
Dominant pathogenic variants in ATAD3A cause defects in neurodevelopment in *Drosophila* and bind Rag GTPases in an ATPase-dependent manner. (a) Schematic of ATAD3A domains and corresponding amino acid positions. Pathogenic mutations are shown in red. (b) The AlphaFold-predicted ATAD3A structure (AF-Q9NVI7-F1) is shown as a green cartoon. ATP (magenta sticks), conserved Walker A (G355, green) and Walker B (E412, gray) motifs are shown as sticks. Lys358 (yellow) from the Walker A motif is positioned to interact with the phosphate groups of ATP. The disease-associated p.R528W substitution was modeled in Coot, and the Trp528 side chain (cyan) is shown at the Sensor 2 position adjacent to the nucleotide-binding site. (c) Table summarizing effects of *dAtad3a* mutant neuronal expression on development in *Drosophila.* (d) Volcano plot representing dAtad3a-V5 immunoprecipitation (IP) followed by mass spectrometry from *Drosophila* larvae expressing dAtad3a-V5 under control of *dAtad3a-T2A-Gal4* (FC>2, p<0.05). Empty vector expression served as a background control. Results are from n=3 independent experiments from ∼250 of 3^rd^ instar larvae. (e) Selected GO terms of significantly enriched proteins identified from dAtad3a-V5 IP mass spectrometry. (f) Table of significantly enriched lysosomal proteins enriched in dAtad3a- V5 IP/MS and human orthologs. (g) IP by HA antibody resin followed by western blot for endogenous ATAD3A from HEK293T cells transfected with HAGST-tagged mTORC1 regulators. (h) IP by HA antibody resin followed by western blot for V5 from HEK293T cells co-transfected with HAGST-RagD and ATAD3A-V5 WT or ATPase mutants.

To assess the functional impacts of pathogenic variants in vivo, we expressed homologous mutations in *Drosophila Atad3a* (*dAtad3a; bor*) including *dAtad3a^R534W^* and *dAtad3a^G361D^* in *Drosophila* neurons. Pan-neuronal expression of either mutation resulted in lethality at pupal stage, indicating that both variants act as dominant-negative mutations (**Figure 1c**). In contrast, expression of the WB motif mutation p.E412Q (*dAtad3a^E418Q^*) did not cause developmental defects (**Figure 1c**). WB mutations slow down or abolish ATP hydrolysis in most AAA+ ATPases but do not affect ATP binding.^43, 44^ Collectively, these findings indicate that ATP binding capacity in ATAD3A is required for normal neurodevelopment.

### ATAD3A interacts with lysosomal proteins, including Rag GTPases

ATAD3A possesses a unique topology allowing for interactions with proteins both inside and outside of the mitochondria.^21, 23^ To shed light on how ATAD3A regulates cellular processes through protein-protein interactions, we performed immunoprecipitation followed by mass spectrometry (IP/MS) from *Drosophila* expressing a C-terminal V5-tagged *dAtad3a* under control of endogenous *dAtad3a* promoter (*dAtad3a-T2A-Gal4>UAS-dAtad3a-V5*; *dAtad3a-T2A- Gal4>UAS-empty* flies as control) (**Figure 1d)**.^26^ Gene Ontology (GO) analysis of the dAtad3a- binding partners indicates an enrichment of proteins involved in biological processes such as oxidative phosphorylation, translation, nucleoside phosphate biosynthetic process, ATP synthesis coupled with electron transport, vesicle-mediated transport, amino acid metabolic process, fatty acid oxidation, establishment of protein localization to organelle, and autophagy (**Figure 1e**). Many interactors with dAtad3a recapitulate the previous studies that demonstrate human ATAD3A interacting with mitochondrial proteins such as those involved in the electron transport chain (ETC) (ND-30/NDUFS3, ND39/NDUFA9, ND-B14.5A/NDUFA7, ox/UQCR10, ND-51/NDUFV1, ND-B14/NDUFA6, Ccdc56/COA3), cristae maintenance (Tom20/TOMM20, Tim23/TIMM23, and Mge/TOMM22), and mitochondrial dynamics (Drp1/DNM1L, Pgam5/PGAM5) (**Figure 1d-e**).^15, 22^ Notably, we found an enrichment of a specific subset of lysosome-localized proteins involved in regulation of mTORC1 signaling (Rheb, Sec13, RagA-B, and RagC-D) as well as many components of the V_1_ sector of the vacuolar ATPase (v-ATPase) (Vha68-1, VhaAC39-1, Vha14-1, Vha36-1, VhaSFD, Vha44, Vha13, Vha68-2, Vha55, and Vha26) (**Figures 1d and 1f**). Given that ATAD3A has been linked to altered mTORC1 signaling and lysosomal expansion,^10, 29, 40^ the identification of Rag GTPases as dAtad3a-interacting proteins was particularly intriguing. Rag complexes recruit mTORC1 and TFEB/TFE3 to lysosomes,^46^ yet a direct link between Rag GTPases and mitochondrial proteins has not been established.

To determine whether the interaction between ATAD3A and Rag GTPases is conserved in human cells, we performed co-IP experiments in HEK293T cells expressing HA-GST tagged mTORC1 regulators, including RagA, RagB, RagC, RagD, and Rheb. Rap2A served as a negative control. Immunoprecipitation followed by Western blotting revealed that endogenous ATAD3A specifically interacts with RagC and RagD, but not with RagA, RagB, Rheb or Rap2A (**Figure 1g**). These results indicate that ATAD3A selectively interacts with RagC/D. Together, our data indicate that ATAD3A physically associates with RagC/D GTPases in flies and humans, raising the possibility that this conserved interaction may contribute to regulation of the mTORC1-TFEB axis.

### Pathogenic ATAD3A mutants exhibit stronger interaction with RagD

ATP-/ADP- bound states regulate substrate engagement of AAA+ proteins.^43^ Because the pathogenic ATAD3A variants are predicted to impair ATPase function (**Figure 1b**), we sought to ask whether these mutations affect the interaction of ATAD3A with its binding partners. To test this, we examined the effects of pathogenic variants p.R528W and p.G355D, as well as ATP-binding-deficient WA mutant p.K358A and ATP-hydrolysis-deficient WB mutant p.E412Q on RagD binding. HEK293T cells were co-transfected with plasmids expressing C-terminal V5- tagged ATAD3A mutants and HA-GST-tagged-RagD followed by immunoprecipitation and Western blotting. Compared to WT ATAD3A, p.R528W, p.G335D, and p.K358A exhibited increased binding to RagD (**Figure 1h**). In contrast, p.E412Q, which is predicted to stabilize an ATP-bound state by preventing ATP hydrolysis, did not show an increase in RagD interaction (**Figure 1h**). Together, these data indicate that impaired ATP-binding, rather than defective ATP hydrolysis, enhances the interaction between ATAD3A and RagD. Consistently, expression of ATP-binding mutants caused lethality and neurodevelopment defects *in vivo*, whereas the ATP hydrolysis mutant did not (**Figure 1c**), linking aberrant ATAD3A-Rag GTPase binding to cellular dysfunction and disease caused by *ATAD3A* mutations.

### ATAD3A p.R528W expression results in activation of cellular stress pathways and CLEAR gene expression

To better understand the molecular consequences of ATAD3A p.R528W expression in human cells, we utilized a lentiviral-mediated overexpression system in the SH-SY5Y neuroblastoma cell line. SH-SY5Y cells were transduced with lentivirus to generate cells expressing an Empty vector control, C-terminally V5 tagged wild-type (WT) ATAD3A, or V5 tagged ATAD3A p.R528W. Following blasticidin selection, cells were grown to confluency and harvested for proteomics and transcriptomics analyses.

Overexpression of ATAD3A WT did not substantially alter the proteome, as only 24 proteins were significantly increased, including ATAD3A, (log_2_FC of >0.263 and p value of <0.05) and only 28 proteins were significantly decreased (log_2_FC of <-0.263 and p value of <0.05) in abundance compared to expression of an empty control (**Extended Figure 1a**). However, comparing cells expressing ATAD3A p.R528W and ATAD3A WT, proteomics analysis revealed a significant decrease (log_2_FC of <-0.263 and p value of <0.05) in abundance of 425 proteins and a significant increase (log_2_FC of >0.263 and p value of <0.05) in 659 proteins (**Figure 2a**). ATAD3A was not identified as a protein with a significant alteration in abundance, suggesting comparable expression of ATAD3A WT and ATAD3A p.R528W in our system. Gene set enrichment analysis using the MSigDB Hallmarks collection^45^ revealed enrichment of the apical junction hallmark among proteins with increased abundance in ATAD3A p.R528W- expressing cells (**Figure 2b**), while less abundant proteins were enriched for hallmarks associated with oxidative phosphorylation, DNA repair and cell cycle regulation, and mTORC1 signaling (**Figure 2b**). These findings are consistent with previous studies showing that *ATAD3A* deficiency in mice or humans reduces oxidative phosphorylation protein expression^15, 16, 19^ Overall, our findings indicate that ATAD3A p.R528W expression results in an impairment of protein expression involved in cellular growth and metabolism, establishing a model for investigating the cellular consequences of ATAD3A p.R528W in human cells.

**Figure 2.**
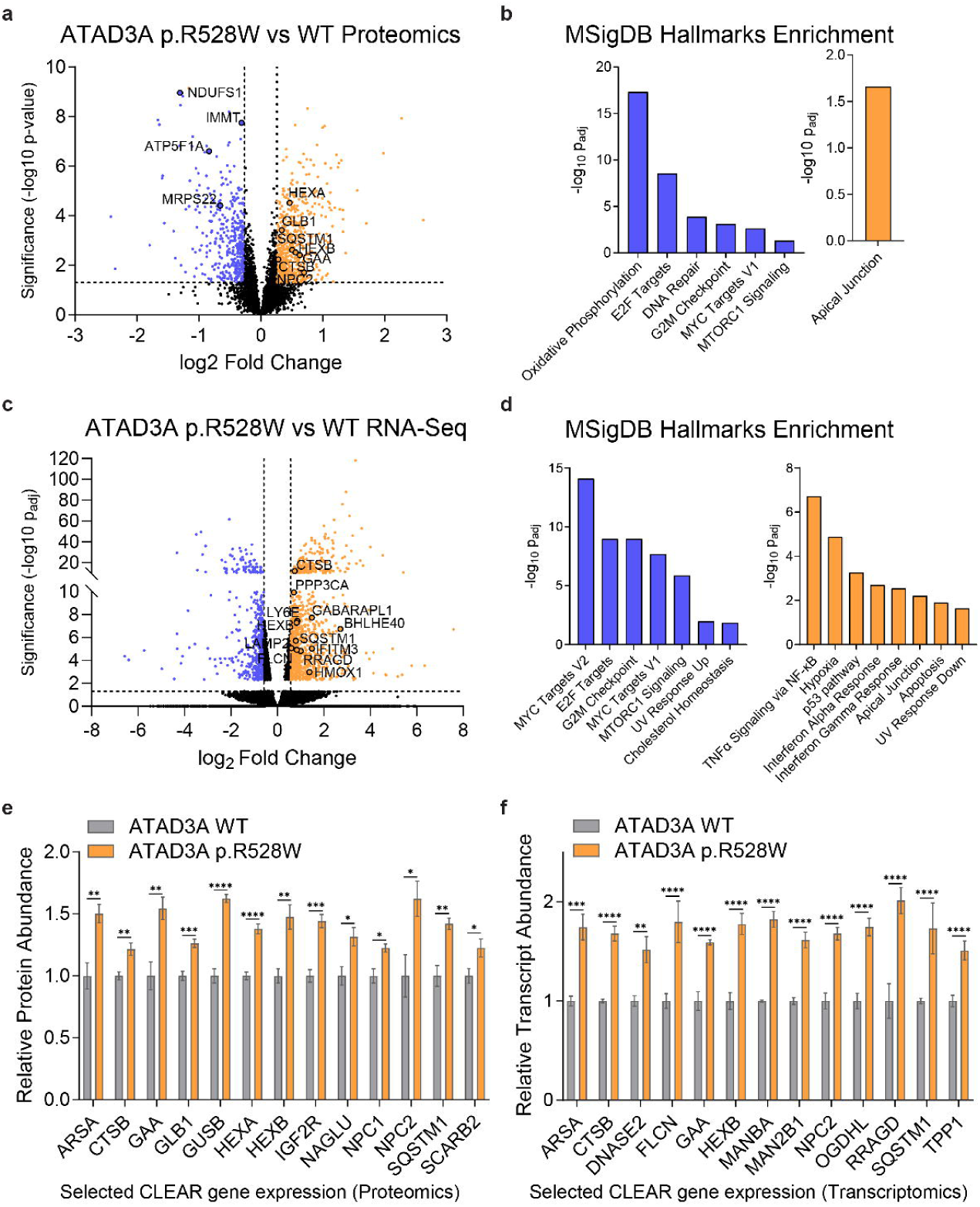
Expression of ATAD3A p.R528W in human cells results in cellular stress including the lysosomal-centric CLEAR gene response. (a) Volcano plot comparing proteomics analysis of SH-SY5Y cells expressing ATAD3A WT-V5 or ATAD3A-V5 p.R528W. Blue represents less abundant (log_2_FC of <-0.263 and p-val <0.05) while orange represents more abundant (log_2_FC of >0.263 and p-val <0.05) proteins in ATAD3A p.R528W-expressing cells. Results are from n=3 independent experiments, with 2 technical replicates run for each. (b) Graphs of significantly enriched MSigDB Hallmarks from proteins more (right) and less (left) abundant in ATAD3A p.R528W-expressing cells compared to those expressing ATAD3A WT. (c) Volcano plot comparing RNA-Seq analysis of SH-SY5Y cells expressing ATAD3A WT-V5 or ATAD3A-V5 p.R528W. Blue represents downregulated genes (log_2_FC of <-0.585 and p_adj_ <0.05) while orange represents upregulated (log_2_FC of >0.585 and p_adj_ <0.05) genes in ATAD3A p.R528W-expressing cells. Results are from n=3 independent replicates. (d) Graphs of significantly enriched MSigDB Hallmarks from transcripts up (right) and downregulated (left) in ATAD3A p.R528W-expressing cells compared to those expressing ATAD3A WT. (e) Selected CLEAR gene products upregulated in ATAD3A p.R528W-expressing cells compared to those expressing ATAD3A WT. Graphs represent mean +/- SEM, n=3 independent experiments with 2 technical replicates run for each. P values were calculated using Student’s t test. *p<0.05, **p < 0.01, ***p < 0.001, ****p < 0.0001 (f) Selected CLEAR genes upregulated in ATAD3A p.R528W-expressing cells compared to those expressing ATAD3A WT. Graphs represent mean +/- SEM, n=3 independent experiments. P_adj_ values were obtained using the Benjamini-Hochberg procedure in DESeq2. **p < 0.01, ***p < 0.001, ****p < 0.0001

Similar to findings at the protein-level, wild-type ATAD3A expression had minimal transcriptomic effects, with RNA-Sequencing (RNA-Seq) identifying only 4 significantly downregulated (log_2_FC of <-0.585 and p_adj_ <0.05) transcripts and ATAD3A as the only significantly upregulated transcript compared with empty control (log_2_FC of >0.585 and p_adj_ <0.05) (**Extended Figure 1b**). Comparing the transcriptome of cells expressing ATAD3A p.R528W and ATAD3A WT, 748 transcripts were downregulated (log_2_FC of <-0.585 and p_adj_ <0.05) while 1378 transcripts were upregulated (log_2_FC of >0.585 and p_adj_ <0.05) (**Figure 2c**). As observed at the protein level, ATAD3A transcript level was not significantly altered. Gene set enrichment analysis of upregulated genes from ATAD3A p.R528W-expressing cells showed an enrichment in genes associated with cellular stress pathways, including metabolic and inflammatory responses, such as hypoxia and interferon responses (**Figure 2d**). ATAD3A p.R528W has been reported to activate interferon responses in human cells^46^, and a hypoxia response can be activated in the context of oxidative phosphorylation defects,^47^ which was observed in the proteome of ATAD3A p.R528W-expressing cells. Downregulated genes in p.R528W-expressing cells include those associated with cell cycle and metabolic pathways (**Figure 2d**). As observed in the proteomic analysis, mTORC1 signaling genes were significantly enriched in downregulated genes from cells expressing ATAD3A p.R528W (**Figure 2d**), consistent with previous studies implicating a role for ATAD3A in mTOR signaling.^29, 40^

Interestingly, these analyses revealed an increase in the levels of proteins and transcripts associated with TFEB activation (**Figure 2e** and **2f**). ^31, 35, 48^ Those include CLEAR genes and their products involved in mTORC1-TFEB regulation (*RRAGD, FLCN)*, lysosomal hydrolases (*CTSB*, *TPP1, DNASE2, MAN2B1, MANBA*, and *GAA, HEXB*), lysosomal cholesterol transport (*NPC2, SCARB2*), and autophagy (*SQSTM1*/p62) (**Figure 2e** and **2f**). We also observed a significant upregulation of *BHLHE40* (FC of 6.55 and p-val of 1.76×10^-7^), and an upregulation trend of *BHLHE41* (FC of 3.04 and p-val 0.21) further suggesting TFEB activation. BHLHE40 and BHLHE41 are established transcriptional regulators during chronic TFEB activation that fine-tune the CLEAR gene transcriptional program.^49^ Together, our proteomic and transcriptomic analyses robustly recapitulate the previously known mitochondrial and metabolic defects associated with ATAD3A dysfunction, while identifying novel mTORC1-TFEB axis associated with *ATAD3A* pathogenic variants.

### Genetic screen in *Drosophila* identifies genes associated with mTORC1-TFEB and autophagy as suppressors of ATAD3A p.R528W

Although our biochemical and cell culture analyses implicate ATAD3A dysfunction in altered mitochondrial metabolism, cellular stress responses, and dysregulation of mTORC1- TFEB signaling, these approaches do not establish which downstream pathways are functionally responsible for the neurodevelopmental defects caused by *ATAD3A* variants in vivo. To address this, we performed a suppression screen by taking advantage of robust developmental lethality caused by neuronal *dAtad3a^R534W^* expression and ample resources of publicly available *Drosophila* RNAi, mutants, and overexpression lines. (**Figure 3a**). As expression of *dAtad3a^R534W^* or *dAtad3a^G361D^* causes comparable neurodevelopmental phenotypes in vivo (**Figure 1c**), we chose to utilize *dAtad3a^R534W^* as a dominant-negative *ATAD3A* disease model for the screen. Among ∼350 RNAi and cDNA lines related to the genes associated with metabolism, growth signaling, mitochondria and autophagy, only twelve genes exhibited robust suppression (i.e., over 30% observed/expected ratio for viability) of the lethality caused by *dAtad3a^R534W^*. (**Figure 3a-b**). The RNAi lines that rescued lethality include genes associated with autophagy (*Autophagy-related 7/Atg7 (44%), Atg8a (48%), Atg1(41%),* and *Rab21 (39%)*), vesicle trafficking (*Rab26 (51%), Rab2 (35%),* and *Rip11 (53%)*), and, and innate immunity (*Sting, 76%*) (**Figure 3b**). Because our previous work demonstrated that expression of *dAtad3a^R534W^* increases autophagy and mitophagy in *Drosophila,*^12^ the identification of autophagy-related genes as genetic suppressors of *dAtad3a^R534W^*-mediated lethality supports aberrant autophagy and mitophagy as key consequences of ATAD3A dysfunction in vivo. In addition, the identification of *Sting* as a suppressor is consistent with the previous observation that patient fibroblasts carrying the *ATAD3A* p.R528W variant exhibit abnormal STING activation,^46^ supporting the link between ATAD3A and innate immune signaling.

**Figure 3.**
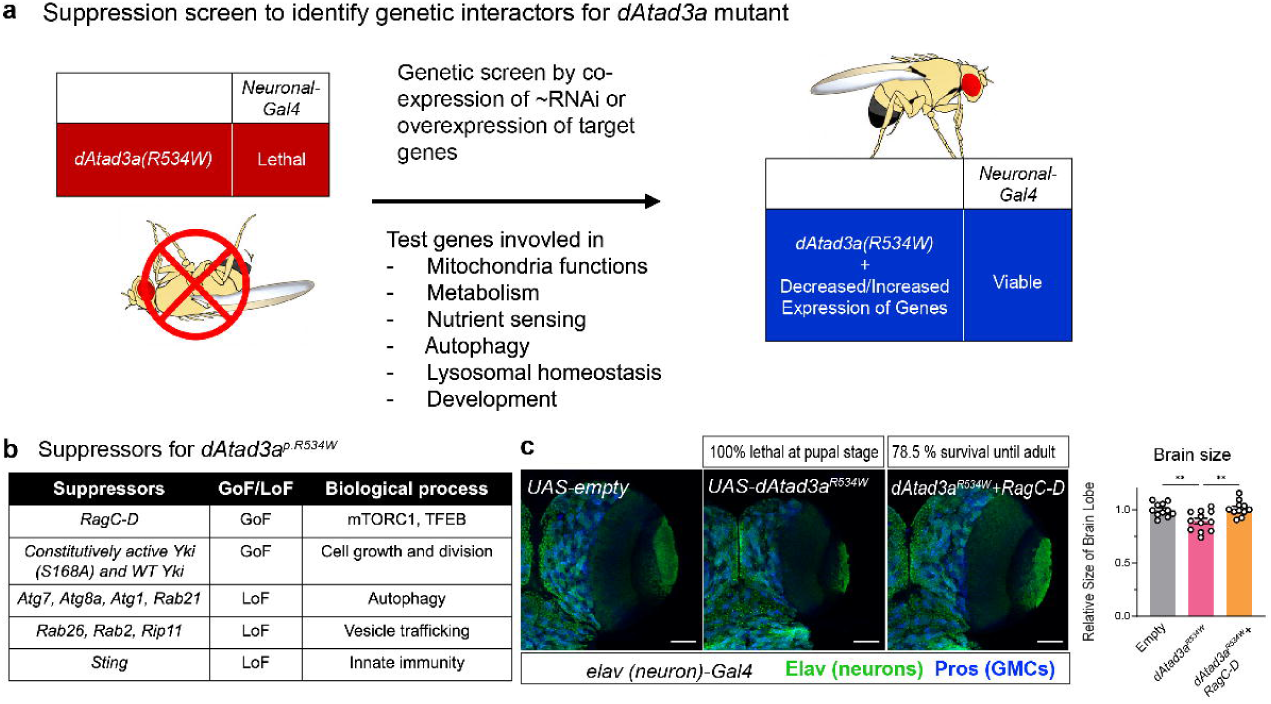
Genetic screen identified RagC-D as a suppressor of dAtad3a p.R534W-indcued neurodevelopmental defects. (a) Schematic of a suppressor screen design to identify genes whose loss or gain rescues developmental lethality caused by neuronal expression of *dAtad3a^p.R534W^* (b) List of the suppressor genes for *dAtad3a^p.R534W^* (c) Confocal micrographs of 3^rd^ instar larvae brains expressing *dAtad3a^p.R534W^* with or without *RagC-D* by neuronal driver (*elav-Gal4*). *UAS-empty* expression serves as control. Elav (green) labels neuron. Pros (blue) labels ganglion mother cells. Scale bar, 50 μm. Relative brain size for each genotype is shown. Error bars indicate SEM. p values were calculated using ordinary one-way ANOVA with Tukey’s multiple- comparisons test. **p < 0.01

The overexpression lines that rescued lethality include *UAS-wild-type yki* and constitutive active *yki (S168A)* (48%), and *UAS-RagC-D* (79%). *Yorkie* (*yki*) is the *Drosophila* homolog of human *YAP1* (*Yes 1 associated transcriptional regulator*) and *RagC-D* is the *Drosophila* homolog of human *RagC* and *RagD* GTPases. Of all lines tested, RagC-D overexpression resulted in the most prominent rescue of *dAtad3a^p.R534W^*-associated lethality and brain development (**Figure 3c**). The RagC/D GTPases play a crucial role in mTORC1 regulation^50–52^ and lysosomal recruitment of TFEB/TFE3.^36, 37, 39^ Functional or genetic deficiency of RagC/D causes impaired mTORC1 regulation and mis-regulation of TFEB/TFE3.^37, 39^ Given the upregulation of TFEB target genes and downregulation mTORC1 signaling genes in human cells expressing ATAD3A p.R528W (**Figure 2**) along with the strong RagC-D rescue of dAtad3a p.R534W-associated lethality (**Figure 3b-c**), we hypothesized that ATAD3A p.R528W may disrupt RagC/D dependent regulation of mTORC1-TFEB axis. Together, our genetic screen revealed that the critical biological consequences resulting from ATAD3A dysfunction include disruption of growth signaling (yki/YAP), vesicle trafficking and autophagy (Atg and Rab genes), innate immunity (Sting), and mTORC1 and TFEB/TFE3 (RagC-D).

### Pathogenic variant ATAD3A p.R528W disrupts the mTOR-TFEB axis

Our findings including the upregulation of CLEAR genes in SH-SY5Y cells expressing ATAD3A p.R528W (**Figure 2**), increased physical interaction between ATAD3A p.R528W and RagD (**Figure 1**), and identification of *RagC-D* as a genetic suppressor of *dAtad3a^R534W^* (**Figure 3**), led us to hypothesize that ATAD3A p.R528W expression disrupts the mTORC1-TFEB axis, leading to aberrant TFEB activation. To test this, we examined alterations in mTORC1 activity and TFEB activation due to ATAD3A p.R528W expression in SH-SY5Y cells. Western blotting revealed a decrease in phospho-S6 ribosomal protein (p-S6) and phospho-4EBP1 (p-4EBP1), indicating a decrease in mTORC1 activation (**Figure 4a-b**). Interestingly, phosphorylation of S6 kinase, another mTORC1 substrate, was not significantly altered. This differential effect suggests that ATAD3A p.R528W may not cause a uniform suppression of mTORC1 activity but rather selectively perturb phosphorylation of a subset of mTORC1 targets. Furthermore, a shift towards a lower molecular weight was observed for TFEB in p.R528W-expressing cells (**Figure 4a-b**), consistent with TFEB dephosphorylation, which promotes TFEB nuclear translocation.^36^ Indeed, cell fractionation followed by Western blotting confirmed an increase in nuclear localization of TFEB, and similarly the regulated transcription factor TFE3, in ATAD3A p.R528W expressing cells compared to cells expressing an Empty control or ATAD3A WT (**Figure 4d**). This is consistent with an upregulation of TFEB target CLEAR genes observed during ATAD3A p.R528W-expression (**Figure 2**). We similarly observed increased nuclear accumulation of Mitf, the *Drosophila* homolog of TFEB/TFE3, when *dAtad3^p.R534W^*was expressed in neuroblasts of *Drosophila* (**Figure 4e**), suggesting this is a conserved consequence of ATAD3A p.R528W in vivo.

**Figure 4.**
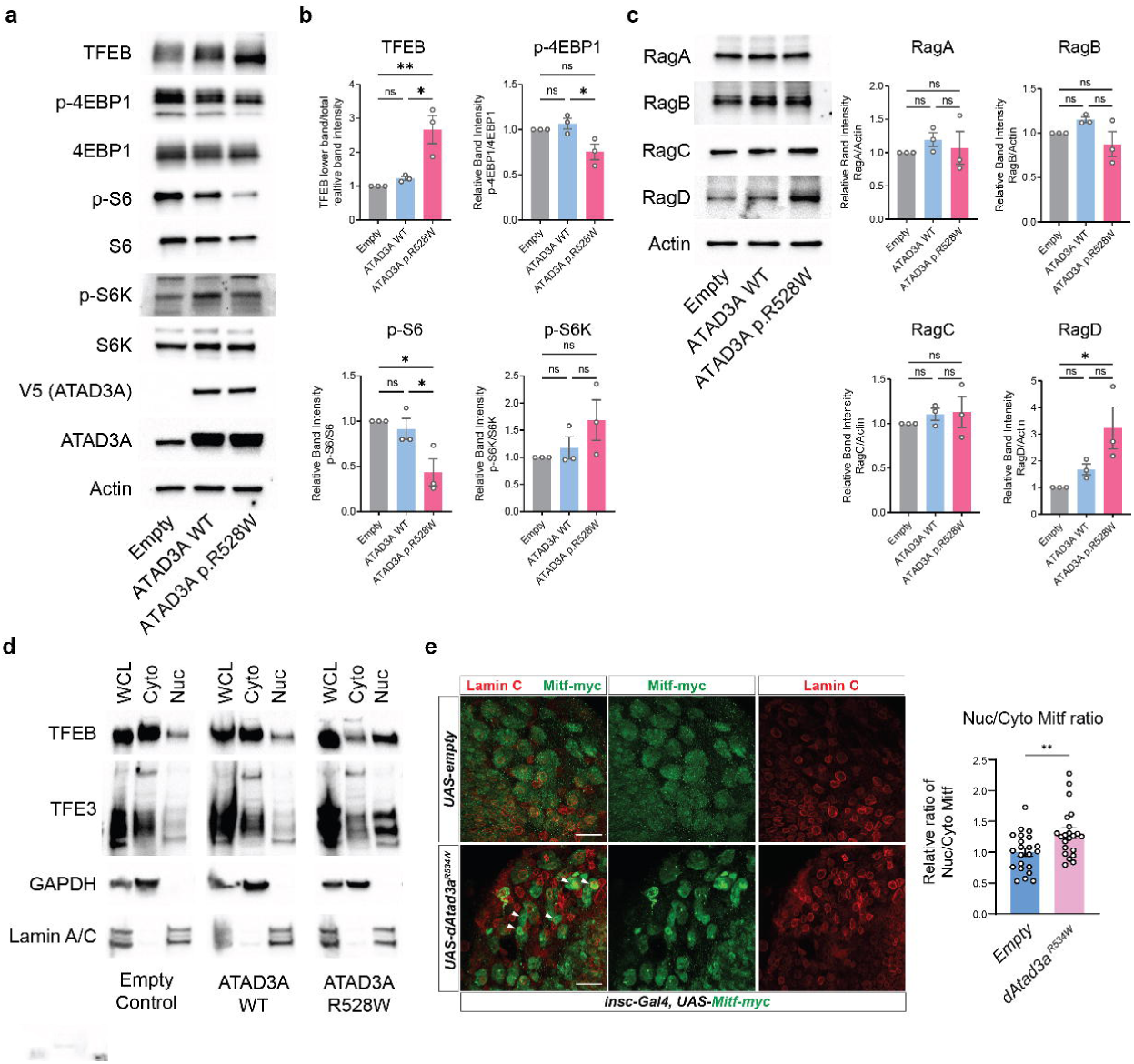
ATAD3A p.R528W expression results in disruption of the mTORC1-TFEB axis. (a) Western blots for mTORC1-TFEB axis proteins from lysates of SH-SY5Y cells expressing Empty control, ATAD3A WT-V5, or ATAD3A p.R528W-V5. Representative images are shown from n=3 independent experiments. (b) Graphs of quantified western blots from (a). Graphs show mean +/- SEM for n=3 independent experiments. P values were calculated using ordinary one-way ANOVA with Tukey’s multiple-comparisons test. ns ≥ 0.05, \**P* < 0.05, and \*\**P* < 0.01. (c) Western blots for Rag GTPases from lysates of SH-SY5Y cells expressing Empty control, ATAD3A WT-V5, or ATAD3A p.R528W-V5 (Left). Representative images are shown from n=3 independent experiments. Graphs of quantified western blots (Right). Graphs show mean +/- SEM for n=3 independent experiments. P values were calculated using ordinary one-way ANOVA with Tukey’s multiple-comparisons test. ns ≥ 0.05 and \**P* < 0.05 (d) Nuclear fractionation from SH-SY5Y cells expressing Empty control, ATAD3A WT-V5, or ATAD3A p.R528W-V5 followed by Western blotting. Representative images of n=2 independent experiments are shown. (e) Confocal micrographs of *Drosophila* brains co-expressing *Mitf-Myc* and Empty control or *dAtad3a^R534W^* in neuroblasts (Left). Brains were stained for Myc (green) and Lamin C (red) (Left). White arrows represent nuclear Mitf. Scale bar, 20 μm. (Right) Quantification of the ratio of nuclear and cytoplasmic Mitf-Myc. Error bars indicate SEM. p values were calculated using Welch’s t test. \*\**P* < 0.01.

Because ATAD3A p.R528W showed enhanced interaction with RagC/D GTPases, we next determined whether this increased binding alters Rag GTPase abundance or stability. Western blot analysis showed that RagA, RagB, and RagC protein levels were not significantly changed in ATAD3A p.R528W-expressing cells (**Figure 4c**), indicating that the p.R528W does not broadly affect Rag GTPase stability or expression. RagD showed a modest but significant increase in protein abundance (**Figure 4c**), consistent with its upregulation at the transcriptional level (**Figure 2f**). Thus, ATAD3A p.R528W does not reduce Rag GTPase protein levels, suggesting that enhanced ATAD3A-Rag binding disrupts the mTORC1-TFEB axis through a mechanism other than altering Rag GTPase stability. Together, these data indicate that ATAD3A p.R528W dysregulates mTORC1 substrate phosphorylation and promotes TFEB nuclear translocation without altering Rag GTPase stability.

### Lyso-IP reveals alterations in the lysosomal proteome due to ATAD3A p.R528W expression

Because pathogenic ATAD3A variants impair lysosomal homeostasis,^10^ and ATAD3A p.R528W enhances interaction with lysosomal Rag proteins (**Figure 1h**), we hypothesized that ATAD3A p.R528W alters the lysosomal protein composition, including Rag GTPases. To test this, we generated SH-SY5Y cells stably expressing TMEM192 3xHA and TMEM192 2xFlag (as a negative control) to allow for immunoprecipitation of intact lysosomes (Lyso-IP) using anti-HA magnetic beads as previously described.^53^ TMEM192 3xHA-expressing cells were then transduced with lentivirus to express an Empty vector control, ATAD3A-V5 WT or ATAD3A-V5 p.R528W. After immunoprecipitation of lysosomes, lysosomal proteins were eluted using NP-40 and subjected to LC/MS. Comparing LC/MS from control TMEM192 2xFlag cells to those expressing TMEM192 3xHA, we observed robust enrichment of lysosomal proteins (**Figure 5a**). LC/MS from TMEM192 3xHA with ATAD3A p.R528W samples revealed significant alterations (FC of >1.2 or <0.833 and p<0.1) of the lysosomal proteome compared to ATAD3A WT-expressing cells (**Figure 5b**).

**Figure 5.**
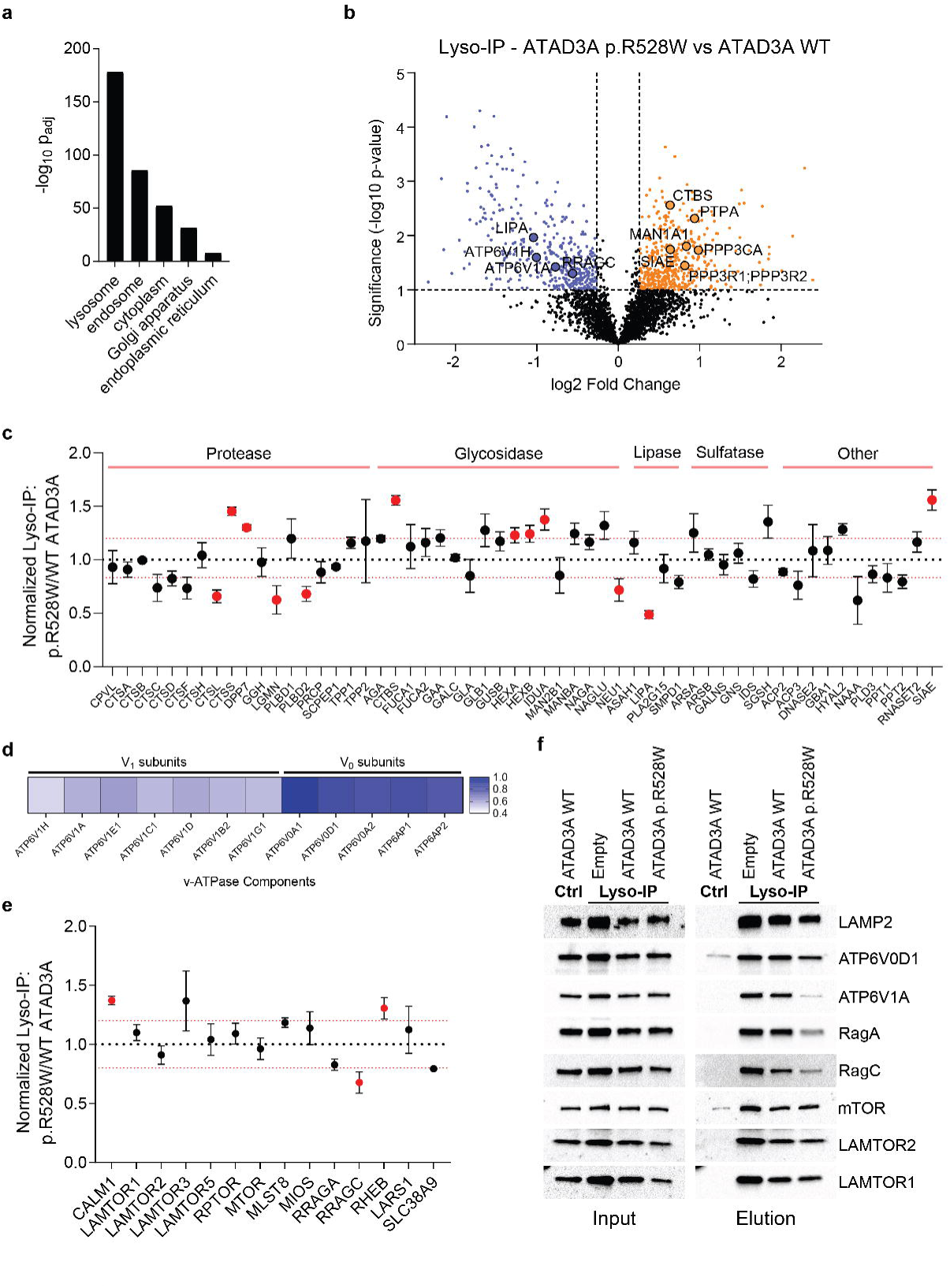
Lyso-IP reveals alterations in the lysosomal proteome due to ATAD3A p.R528W expression. (a) GO Cellular Component results from significantly enriched (p<0.05) proteins after anti-HA pulldown from cells expressing a TMEM192 3xHA versus TMEM192 2xFlag control. Significantly enriched proteins are from n=3 independent experiments. (b) Volcano plot from LC/MS of anti- HA elution fractions from Lyso-IP (cells expressing TMEM192 3xHA) of ATAD3A p.R528W vs ATAD3A WT-expressing cells. Results from n=3 independent experiments, significance is FC of >1.2 or <0.833 and p<0.1. Orange represents proteins enriched in ATAD3A p.R528W Lyso-IP elutions while blue are those enriched in ATAD3A WT Lyso-IP elutions. (c) Scatterplot of lysosomal hydrolase abundance by LC/MS from elution fractions of Lyso-IP. Dots show mean ratio +/- SEM for ATAD3A p.R528W/ATAD3A WT-expressing cell Lyso-IP. n=3 independent experiments. P values were calculated using Student’s t-test. Significantly altered (FC of >1.2 or <0.833 and p<0.1) proteins are labeled in red (d) Heat map of V_1_ (left) and V_O_ (right) v-ATPase component Lyso-IP elution mean ratio of ATAD3A p.R528W/ATAD3A WT cells. N=3 independent experiments (e) Scatterplot of mTORC1-associated protein abundance by LC/MS from elution fractions of Lyso-IP. Dots show mean ratio +/- SEM for ATAD3A p.R528W/ATAD3A WT expressing cell Lyso-IP. N=3 independent experiments. Significantly altered (FC of >1.2 or <0.833 and p<0.1) proteins are labeled in red. p values were calculated using Student’s t-test. (f) Western blot for LAMP2 (lysosomal marker), v-ATPase, or mTORC1-related proteins from input (left) and elution (right) fractions of Lyso-IP from Control (TMEM192 2xFlag + ATAD3A WT), or Lyso-IP (TMEM192 3xHA) cells expressing an Empty control, ATAD3A WT, or ATAD3A p.R528W. Representative images of n=3 independent experiments are shown.

We observed altered protein levels of a subset of lysosomal hydrolases in the cells expressing ATAD3A p.R528W compared to those expressing WT ATAD3A (**Figure 5b and 5c**). CTSS (cathepsin S), DPP7 (dipeptidyl-peptidase 7), CTBS (chitobiase), HexA and HexB (beta-hexosaminidase subunit A and B), IDUA (alpha-L-iduronidase), and SIAE (sialic acid acetylesterase) were enriched in the lysosomes of the cells expressing p.R528W, whereas CTSL (Cathepsin L), LGMN (legumain), PLBD2 (phospholipase B domain containing 2), NEU1 (neuraminidase 1), and LIPA (lysosomal acid lipase) were significantly reduced (**Figure 5c**). Of particular note, LAL/LIPA, which catabolizes cholesteryl esters and triglycerides to generate free fatty acids and free cholesterol,^54, 55^ was significantly reduced in lysosomes of ATAD3A R528W- expressing cells, consistent with previous observation in which lysosomal cholesterol is elevated in patient cells carrying pathogenic *ATAD3A* variants.^10^ Interestingly, lysosomes isolated from ATAD3A p.R528W-expressing cells showed marked depletion of all v-ATPase V_1_ subunits (ATP6V1A, ATP6V1B2, ATP6V1C1, ATP6V1D, ATP6V1E1, ATP6V1H, ATP6V1G1) (**Figure 5d**). Despite depletion of all the V_1_ subunits, the membrane-embedded V_O_ subunits are overall preserved in lysosomes of the cells expressing p.R528W (**Figure 5d**), suggesting that ATAD3A p.R528W-expression impairs the assembly or trafficking of V_1_ subunits to the lysosomal V_O_ subunits.

Notably, Lyso-IP revealed altered lysosomal association of mTORC1-TFEB regulators in ATAD3A p.R528W-expressing cells. These include enrichment of phosphatases that target TFEB (PPP3CA, PPP3R1, PPP3R2, and PTPA) (**Figure 5b)** and reduced Rag GTPase RagC and RagA (**Figure 5e**). We observed a significant decrease in RagC abundance (FC of 0.678 and p-val: 0.049) and a notable decrease in RagA (FC of 0.828, p-val: 0.196) from lysosomes of ATAD3A p.R528W-expressing cells, while RagB and RagD were not successfully detected by LC/MS. In contrast, Lyso-IP did not reveal significant changes in other mTORC1-related proteins such as mTOR, LAMTOR1, LAMTOR 2, LAMTOR 3, LAMTOR 5, and RPTOR in ATAD3A p.R528W-expressing cells (**Figure 5e**).

To validate a subset of these findings including altered lysosomal Rag GTPases and v-ATPase components, we performed Western blotting from input and elution fractions of Lyso-IP samples. By assessing the levels for the V_O_ subunit ATP6V0D1 and V_1_ subunit ATP6V1A, we found no marked decrease ATP6V0D1, while a substantial decrease in ATP6V1A in Lyso-IP elutions from cells expressing ATAD3A p. R528W (**Figure 5f**). Western blotting also confirmed a decrease in RagA and RagC protein levels on lysosomes of ATAD3A p.R528W-expressing cells compared to those of the cells expressing Empty controls or ATAD3A WT (**Figure 5f**). Because the Ragulator complex anchors Rag GTPases to the lysosomal surface,^38^ we assessed the localization of Ragulator complex proteins, including p18 (LAMTOR1) and p14 (LAMTOR2). Validating our findings by LC/MS, Western blot analysis showed that p18 and p14 were not decreased in the elution fractions from ATAD3A p.R528W-expressing cells (**Figure 5f**). Therefore, decreased lysosomal-localization of Rag GTPases is likely not due to altered lysosomal localization of the Ragulator complex. Western blotting also showed that mTOR abundance at lysosomes was unchanged in ATAD3A p.R528W-expressing cells (**Figure 5f**), suggesting that ATAD3A p.R528W preferentially disrupts lysosomal Rag GTPase recruitment without significantly affecting mTOR localization. Together, these data support a model in which ATAD3A p.R528W reduces lysosomal Rag GTPase recruitment, thereby disrupting the TFEB/TFE3 axis.

### RagC-D expression rescues development defects and impaired lysosomal homeostasis caused by ATAD3A mutations

Given that Lyso-IP showed depletion of not only RagC but also RagA in cells expressing ATAD3A p.R528W (**Figure 5f**), we sought to determine whether expression of *RagA-B*, the *Drosophila* homolog of human *RagA* and *RagB*, could rescue the lethality caused by *dAtad3a^p.R534W^*, as observed with *RagC-D* (**Figure 3c**). To this end, we generated flies carrying *UAS-wild-type RagA-B* inserted at the same genomic locus (attP2) as *UAS-RagC-D*.^56^ We found that the flies expressing *dAtad3a^p.R534W^*together with *RagA-B* exhibit 53% viability to the adult stage, rescuing lethality to a lesser extent than flies co-expressing *dAtad3a^p.R534W^*and *RagC-D* (79%) (**Figure 6a**). Furthermore, over 50% of the adult flies expressing *dAtad3a^p.R534W^*and *RagA-B* died within 3 days, whereas flies expressing *dAtad3a^p.R534W^* and *RagC-D* exhibited a similar lifespan to controls (*UAS-empty* and *UAS- dAtad3a^WT^*) (**Figure 6b**). These data indicate that *RagA-B* is also a genetic suppressor of *dAtad3a^p.R534W^*, while *RagC-D* is a more critical downstream effector of *dAtad3a^p.R534W^* in vivo.

**Figure 6.**
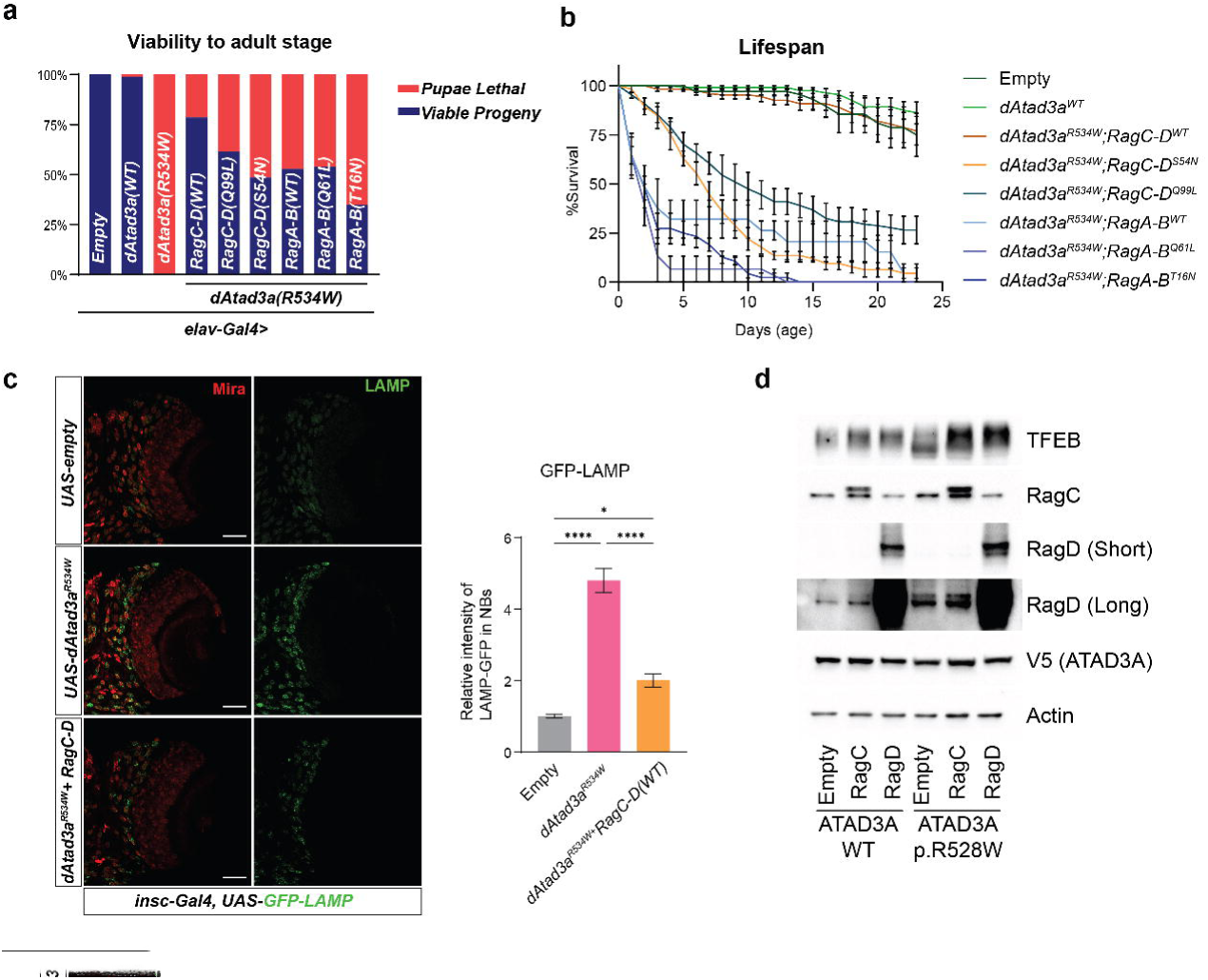
Expression of RagC-D rescues neurodevelopmental and Mitf/TFEB-dependent lysosomal expansion. (a-b) Graph of viability (a) and lifespan (b) for *Drosophila* expressing *Atad3a^p.R534W^*with or without wildtype or GTP or GDP-locked RagC-D or RagA-B in neuronal cells. (c) Confocal micrographs of *Drosophila* brains co-expressing LAMP-GFP lysosomal reporter along with controls, *Atad3a^p.R534W^* with or without *RagC-D* in neuroblasts (left). Mira (red) labels neuroblasts. Scale bar, 50 μm. Quantification of LAMP-GFP intensity (right). Graphs represent mean and error bars indicate SEM. p values were calculated using one-way ANOVA. *p < 0.05, ****p < 0.0001 (d) Western blotting for lysates of SH-SY5Y cells co-expressing WT or p.R528W ATAD3A along with Empty control, FLAG-RagC, or FLAG-RagD to examine TFEB molecular weight shift. Images are representative of N=2 independent experiments.

The nucleotide-bound state of the Rag GTPases regulates mTORC1 complex and TFEB/TFE3 recruitment to the lysosomal surface.^36, 51, 52, 57^ GTP-bound RagA/B promotes recruitment of the mTORC1 complex through Raptor, whereas GDP-bound RagC-D is required for engagement of TFEB/TFE3 on lysosomes.^36, 37, 52, 57^ We therefore asked whether the GTP- or GDP-bound states of Rag GTPases influence rescue phenotypes when co-expressed neuronally with *dAtad3a^p.R534W^*. To test this, we generated flies carrying *UAS-RagC-D^S54N^*(GDP- bound, constitutively active), *UAS-RagC-D^Q99L^* (GTP-bound, inactive), *UAS-RagA-B^Q61L^* (GTP- bound, constitutively active), and *UAS-RagA-B^T16N^* (GDP-bound, inactive) inserted at the same genomic locus for *wild-type UAS-RagA-B* and *UAS-RagC-D*.^51, 58, 59^ We found that expression of both *RagC-D^S54N^* and *RagC-D^Q99L^* restored pupal-to-adult viability to 50–60%, which is lower than wild-type *RagC-D* expression (79%) (**Figure 6a**). Similarly, expression of *RagA-B^Q61L^* or *RagA-B^T16N^*produced rescue effects comparable to, or weaker than, those shown in wild-type *UAS-RagA-B* (**Figure 6a**). However, none of the GTP- or GDP-locked Rag GTPase variants restored lifespan to the extent observed with wild-type *RagC-D* (**Figure 6b**). Thus, these data indicate that impaired regulation of Rag nucleotide state is unlikely to be the primary downstream defect caused by *dAtad3a^p.R534W^*. Rather, a deficiency of wild-type RagC-D appears to be the primary consequence of pathogenic *dAtad3a^p.R534W^*expression in vivo.

Our previous studies showed that expression of *dAtad3a^p.R472C^*, corresponding to the human ATAD3A p.R466C variant, causes lysosomal expansion in neuroblasts (NBs) of the *Drosophila* brain.^10^ Consistent with this finding, expression of *dAtad3a^p.R534W^* in neuronal cells, including NBs increased the lysosomal marker GFP-LAMP1 and enlarged lysosomal volume (**Figure 6c**). Because ATAD3A p.R528W and its *Drosophila* counterpart *dAtad3a^p.R534W^* promote nuclear localization of TFEB/TFE3 in SH-SY5Y cells and Mitf in *Drosophila* neuroblasts (**Figure 4d-e**), these findings suggest that aberrant activation of the TFEB/TFE3/Mitf transcriptional program drives lysosomal expansion. Given the established role of RagC-D in regulating TFEB/TFE3 subcellular localization and TFEB/Mitf as regulators of lysosomal biogenesis,^36, 37^ we hypothesized that restoring *RagC-D* expression would suppress the lysosomal phenotypes caused by *dAtad3a^p.R534W^*. Indeed, wild-type *RagC-D* expression substantially suppressed the lysosomal expansion caused by *dAtad3a^p.R534W^*(**Figure 6c**). Similarly, co-expression of RagC or RagD with ATAD3A p.R528W in SH-SY5Y cells restored the TFEB mobility shift toward the pattern observed in control cells (**Figure 6d**), consistent with restored TFEB phosphorylation and cytoplasmic retention. Together, these data indicate that RagC/D expression suppresses aberrant TFEB/Mitf activation and lysosomal expansion caused by pathogenic ATAD3A variants. Collectively, our in vivo and in vitro data indicate that ATAD3A p.R528W and its *Drosophila* counterpart *dAtad3a^p.R534W^* disrupt RagC/D-dependent regulation of TFEB-mediated lysosomal homeostasis and underscore RagC/D is a critical downstream effector of ATAD3A pathogenic mutations.

### Pathogenic effects of p.R528W are mediated through the N-terminal coiled-coil domain of ATAD3A

Because ATAD3A p.R528W exhibits increased binding to Rag GTPases and disrupts their lysosomal localization, we next sought to define the domains within ATAD3A that mediate binding to RagC/D. To this end, we performed domain-mapping experiments by generating N-terminal deletions of the first 80 residues (ΔN80), the coiled-coil domain (ΔCC), both the first 80 residues and coiled-coil domain (ΔN80-CC), and the first 245 amino acids of ATAD3A (ΔN245), encompassing almost all of the protein outside of the transmembrane domain (**Figure 7a**). Co- transfections of full length or ATAD3A N-terminal truncations with HA-GST RagD followed by anti-HA immunoprecipitation and Western blotting revealed ATAD3A binding of RagD by only full length or ΔN80 constructs (**Figure 7b**). Therefore, loss of binding occurred in any truncations encompassing the ATAD3A coiled-coil domain, indicating that the coiled-coil domain is required for ATAD3A to interact with RagD.

**Figure 7.**
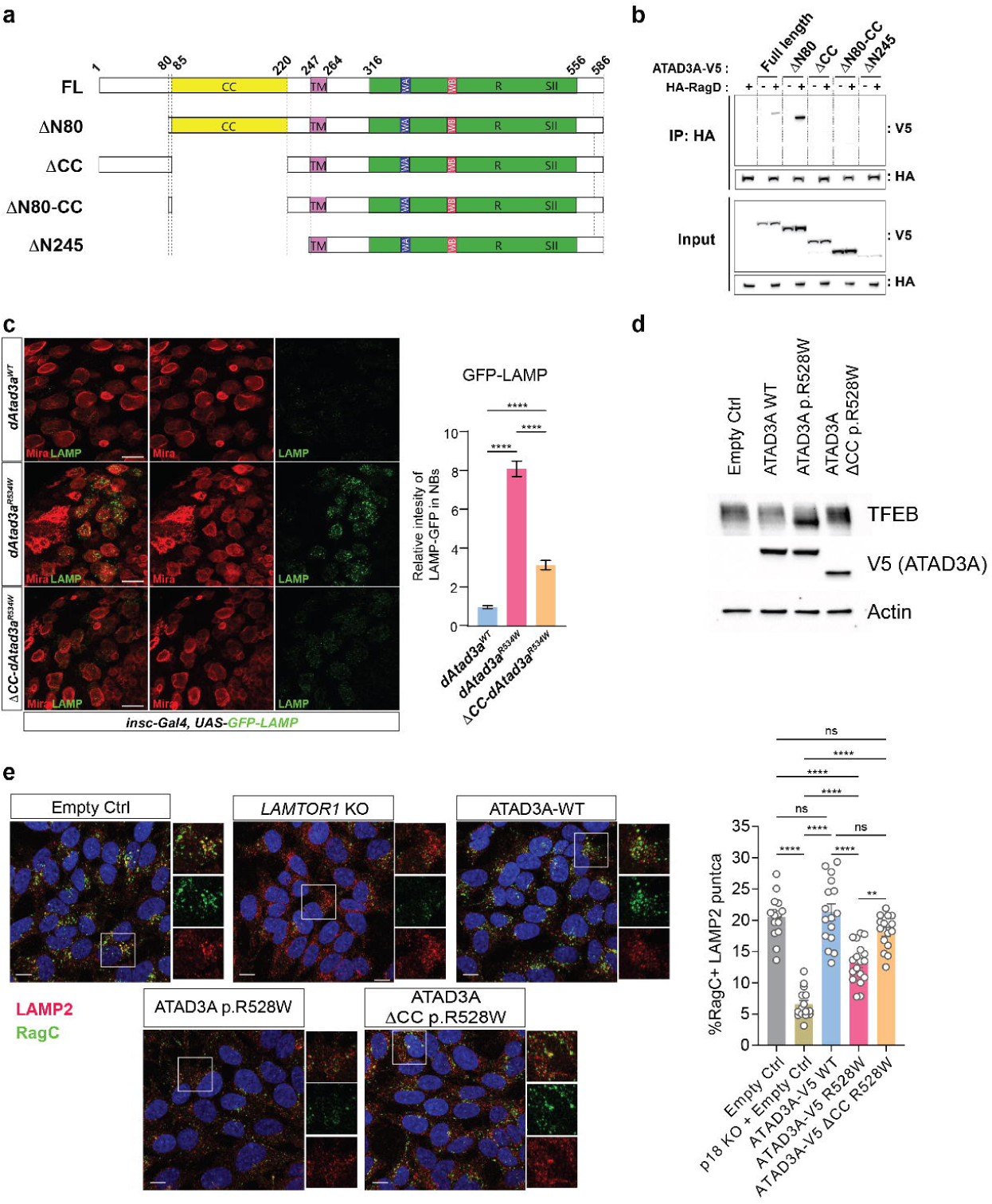
Pathogenic effects of p.R528W are mediated through the N-terminal coiled-coil domain of ATAD3A. (a) Schematic of truncated ATAD3A proteins used in domain mapping experiments. (b) IP followed by Western blotting for truncated ATAD3A proteins co-expressed with HAGST-tagged RagD in HEK293T cells. (c) Confocal images of *Drosophila* larval brains co-expressing LAMP- GFP along with controls (empty), *Atad3a^p.R534W^*, or *ΔCC-Atad3a^p.R534W^* in neuroblasts (left). Mira (red) labels neuroblasts. Scale bar, 20 μm. Quantification of LAMP-GFP intensity in neuroblasts (right). Error bars indicate SEM. p values were calculated using one-way ANOVA. ****p < 0.0001 (d) Western blot for TFEB molecular weight shift in lysates from SH-SY5Y cells expressing Empty control, ATAD3A WT, ATAD3A p.R528W, or ATAD3A ΔCC p.R528W. Images are representative of n=2 independent experiments. (e) Confocal images (left) of SH-SY5Y cells expressing controls or ATAD3A mutants stained for RagC (green) and LAMP2 (red). Scale bar, 10 μm. Quantification of RagC-positive LAMP2 puncta (right). Graphs represent mean +/- SEM from at least 15 images. p values were calculated using ordinary one-way ANOVA with Tukey’s multiple-comparisons test. ns ≥ 0.05, \*\**P* < 0.01, and \*\*\*\**P* < 0.0001.

Because pathogenic ATAD3A variants, including p.R528W, show enhanced RagD binding that requires the coiled-coil domain, we hypothesized that p.R528W pathogenicity is mediated by the ATAD3A coiled-coil domain-RagD association. Therefore, deletion of the coiled-coil domain may suppress the lysosomal phenotypes caused by ATAD3A p.R528W. To test this, we generated flies carrying *UAS-dAtad3a^p.R534W^(ΔCC)* together with *UAS-dAtad3a^WT^(ΔCC).* Flies expressing *dAtad3a^WT^ (ΔCC)* in neurons show no developmental lethality, similar to those in *dAtad3a^WT^*expression (**Extended Figure 2**). In contrast to the lethality caused by full-length *dAtad3a^p.R534W^*, most flies expressing *dAtad3a^p.R534W^(ΔCC)* in neurons remained viable, indicating that the pathogenic effects of *dAtad3a^p.R534W^* on development requires the coiled-coil domain (**Extended Figure 2**). We next examined whether this domain also contributes to lysosomal expansion and found that the *dAtad3a^p.R534W^(ΔCC)* resulted in significantly less GFP- LAMP1 accumulation compared to full-length *dAtad3a^p.R534W^*(**Figure 7c**). Together, these data indicate that the coiled-coil domain is necessary for the pathogenic effects of the *ATAD3A* mutations on both neurodevelopmental defects and lysosomal expansion.

We next tested whether the coiled-coil domain mediates TFEB activation in the human cells expressing ATAD3A p.R528W. Using the lentiviral system previously described, we expressed controls (ATAD3A WT), ATAD3A p.R528W, or ATAD3A ΔCC p.R528W and assessed TFEB mobility shift by Western blotting as a proxy of activation state. Deletion of the coiled-coil domain in ATAD3A p.R528W did not affect the protein stability or mitochondrial localization (**Figure 7d** and **Extended Figure 3b**). Interestingly, while ATAD3A p.R528W expression results in TFEB lower molecular weight shift, ATAD3A ΔCC p.R528W-expressing cells have a TFEB molecular weight pattern similar to that of ATAD3A WT-expressing cells (**Figure 7d**), indicating that TFEB activation caused by ATAD3A p.R528W requires the coiled-coil domain.

Because RagC abundance was reduced on lysosomes from ATAD3A p.R528W- expressing cells (**Figure 5**), and ATAD3A binding of Rag GTPases was mediated by the coiled-coil domain (**Figure 6Aa**), we hypothesized that this domain contributes to RagC mislocalization in cells expressing ATAD3A p.R528W. To further define whether the coiled-coil domain affects lysosomal localization of Rag GTPases and/or mTOR under conditions of ATAD3A dysfunction, we performed immunocytochemistry analysis for RagC together with lysosomal marker LAMP2. As a control for disrupted RagC localization, we also generated SH-SY5Y cells lacking p18 (LAMTOR1) which is required for lysosomal localization of Rag GTPases (**Extended Figure 3a**).^38, 60^ Consistent with its known molecular function, cells lacking p18 (*LAMTOR1* KO) exhibited reduced numbers of RagC-positive LAMP2 puncta compared to cells expressing Empty, or ATAD3A WT (**Figure 7e**). *p18/LAMTOR1* KO also exhibited a TFEB mobility shift in SDS-PAGE, linking Rag GTPase mislocalization to TFEB activation (**Extended Figure 3a**). Cells expressing ATAD3A p.R528W showed a significant reduction in RagC-positive LAMP2 puncta, indicating impaired lysosomal localization of RagC (**Figure 7e**). In contrast, ATAD3A

ΔCC p.R528W cells exhibited comparable numbers of lysosomal-localized RagC puncta to those in Empty control, and ATAD3A WT cells, suggesting that the coiled-coil domain of ATAD3A is required for p.R528W-mediated reduction in lysosomal RagC localization. Interestingly, co-staining for mTOR and LAMP2 revealed no significant decrease in mTOR lysosomal localization in ATAD3A p.R528W (**Extended Figure 3c**). This is consistent with Lyso-IP followed by LC/MS which demonstrated no significant alteration in mTOR abundance from ATAD3A p.R528W- expressing lysosomes (**Figure 5f**). Cells lacking p18 exhibited statistically a significant decrease in mTOR lysosomal localization, albeit with less effects on mTOR and LAMP2 co-localization compared to RagC localization on lysosomes (**Extended Figure 3c** and **Figure 7e**). Collectively, these findings reveal that the coiled-coil domain of ATAD3A is required for pathogenic mislocalization of Rag GTPases, TFEB activation, and neurodevelopmental defects.

### ATAD3A p.R528W pathogenic phenotypes are rescued by Mitf knockdown

Pathogenic ATAD3A mutations induce excessive lysosomal biogenesis, likely through aberrant activation of *Mitf* (**Figure 4e**). Our previous work showed that these expanded lysosomes are functionally impaired, as evidenced by the accumulation of intra-lysosomal membrane whorls.^10^ Given that aberrant TFEB/TFE3 activation and lysosomal expansion have been implicated in neurodevelopmental disorders,^61, 62^ we hypothesized that expansion of dysfunctional lysosomes contributes to the neurodevelopmental defects caused by ATAD3A mutations. To test this hypothesis, we reduced lysosomal biogenesis through RNAi-mediated knockdown of *Mitf*. Consistent with the known role of *Mitf* in lysosomal expansion,^63^ knockdown of *Mitf* significantly suppressed the lysosomal expansion caused by *dAtad3a^p.R534W^* (**Figure 8a**). Importantly, reducing lysosomal biogenesis through *Mitf* knockdown significantly restored brain size and rescued the developmental lethality caused by neuronal expression of dAtad3a^p.R534W^ (**Figure 8b-c**). Together, these results indicate that aberrant TFEB/Mitf-mediated lysosomal expansion contributes to *ATAD3A* mutation-induced neurodevelopmental defects and that limiting this expansion mitigates pathogenic consequences of ATAD3A dysfunction.

**Figure 8.**
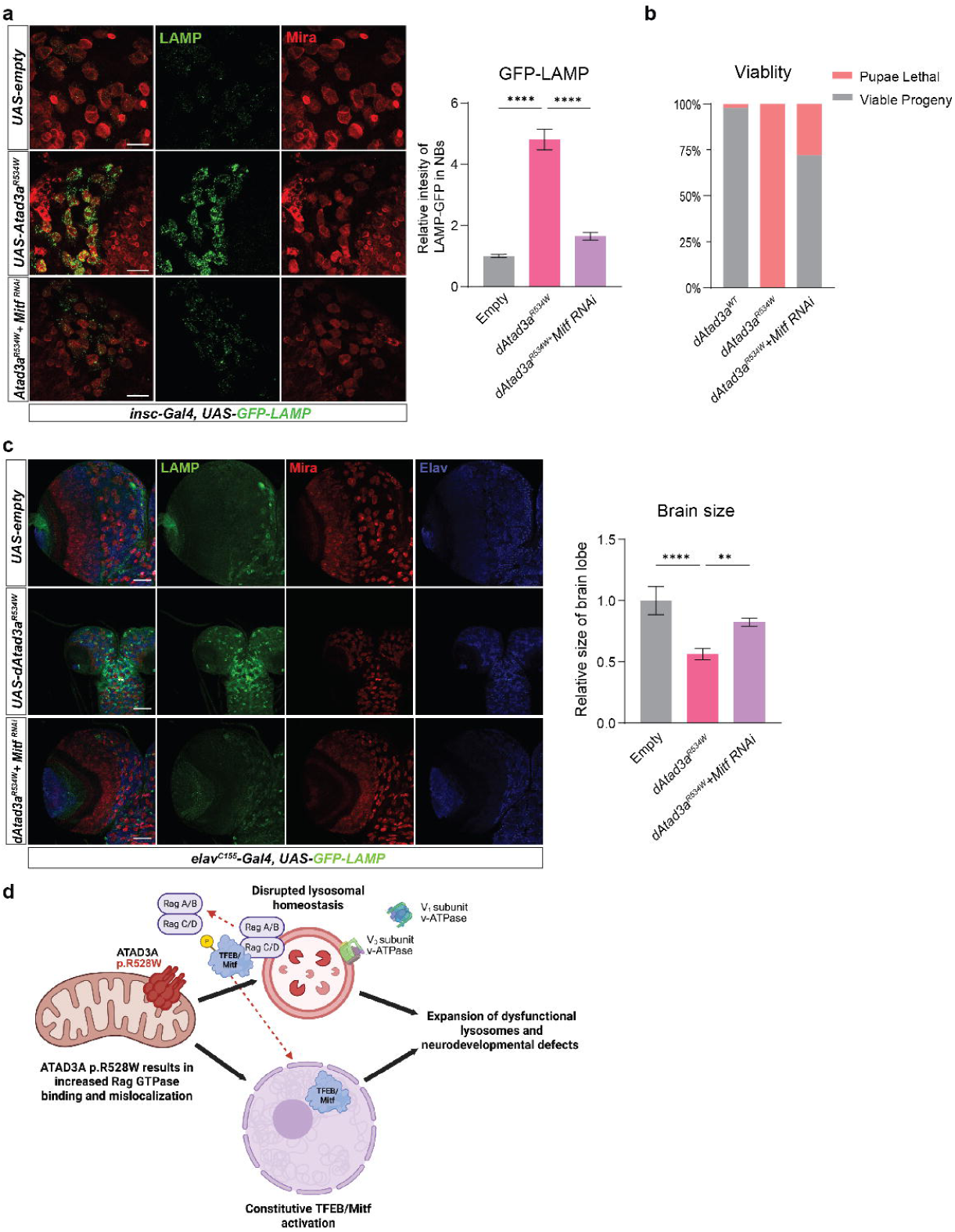
Mitf knockdown rescues dAtad3a p.R534W-dependent lysosomal expansion and neurodevelopmental defects. (a) Confocal micrographs of *Drosophila* brains co-expressing LAMP1-GFP along with controls, *Atad3a^p.R534W^*with or without *Mitf* RNAi in neuroblasts (left). Mira (red) labels neuroblasts. Scale bar, 20 μm. Quantification of LAMP1-GFP intensity (right). Error bars indicate SEM. p values were calculated using one-way ANOVA. ****p < 0.0001 (b) Graph of viability for *Drosophila* expressing *Atad3a^WT^*, or *Atad3a^p.R534W^*with or without *Mitf* RNAi in neuronal cells (*elav-Gal4*). (c) Confocal micrographs of *Drosophila* brains co-expressing LAMP1-GFP along with controls, *Atad3a^p.R534W^* with or without *Mitf* RNAi in neuronal cells (left). Mira (red) labels neuroblasts. Elav (blue) labels neurons. Scale bar, 50 μm. Quantification of brain sizes. Error bars indicate SEM. p values were calculated using one-way ANOVA. **p < 0.01, ****p < 0.0001 (d) Model of ATAD3A p.R528W-mediated dysregulation of the RagC/D-TFEB axis, disruption of lysosomal homeostasis and function, and neurodevelopmental defects. Graphic was created using BioRender.

## DISCUSSION

Mitochondrial and lysosomal dysfunction are increasingly recognized as interdependent contributors to human disease, in which primary impairment of one organelle can impose collateral damage on the other and accelerate cellular pathology.^1–3^ For example, loss of mitochondrial proteins such as *Pink1* or *Opa1* impairs lysosomal activity through reactive oxygen species (ROS) in mouse embryonic fibroblasts,^4^ whereas impaired mitochondrial respiration in *Tfam*-deficient mouse T cells disrupts lysosomal function through altered NAD^+^/NADH balance. ^64^ These studies largely implicate diffusible stress signals, including ROS and metabolites, as mediators of mitochondria-to-lysosome dysfunction. Here, we identify a distinct mechanism in which pathogenic ATAD3A, a mitochondrial membrane protein with a unique topology that enables extramitochondrial interactions, disrupts lysosomal mTORC1- TFEB signaling by impairing RagC/D-dependent regulation and lysosomal localization. This mechanism is supported by our genetic screen, which identified *RagC/D*, a lysosomal protein, rather than canonical mitochondrial proteins, as one of the strongest suppressors of ATAD3A pathogenic phenotypes. Remarkably, ectopic expression of *RagC-D* or *TFEB*/*Mitf* knockdown restored lysosomal abundance and rescued neurodevelopmental defects caused by the pathogenic *ATAD3A* variant in *Drosophila*.

Multiple lines of genetic, biochemical, and cell biological evidence identify RagC/D as a critical downstream mediator of ATAD3A pathogenicity both in vitro and in vivo. IP/MS analysis identified RagC-D, together with RagA-B, among candidate dAtad3a-interacting proteins. Consistent with this finding, co-IP experiments in HEK293T cells confirmed that human ATAD3A interacts with RagC and RagD, and that pathogenic variants, including p.R528W, enhance this association. Lyso-IP followed by Western blotting in SH-SY5Y cells further showed that ATAD3A p.R528W reduces lysosomal levels of Rag GTPases, including RagC, and immunocytochemistry confirmed a substantial decrease in RagC colocalization with LAMP2- positive lysosomes. Importantly, our *Drosophila* genetic screen identified *RagC-D* as a strong suppressor of both neurodevelopmental defects and lysosomal expansion caused by pathogenic *dAtad3a^R534W^*. In parallel, overexpression of human RagC or RagD restored TFEB phosphorylation in SH-SY5Y cells expressing ATAD3A p.R528W, consistent with direct role of RagC/D in TFEB/TFE3 recruitment to lysosomes and their cytoplasmic retention.^36, 37, 39, 65^ Together, these complementary findings indicate that impaired lysosomal RagC/D function is a central biological consequence of pathogenic ATAD3A mutations in both flies and human cells.

Our Lyso-IP experiments further revealed reduced lysosomal abundance of RagA in SH- SY5Y cells expressing ATAD3A p.R528W, suggesting that impaired lysosomal recruitment of RagA/B may also contribute to ATAD3A p.R528W-associated pathology. Consistent with this possibility, ectopic expression of *RagA-B* partially rescued *dAtad3a^p.R534W^*-associated developmental lethality and adult lifespan in *Drosophila*, although this suppression was less robust than that achieved by *RagC-D*. Interestingly, loss of *RagA/B* has also been reported to promote aberrant nuclear localization of TFEB in multiple tissues.^66, 67^ This phenotype may arise from destabilization of RagC/D proteins in the absence of RagA/B or from the cooperative requirement of RagA/B–RagC/D heterodimers for efficient TFEB engagement at lysosomes.^65, 68, 69^ Notably, heart-specific deletion of both *RagA* and *RagB* causes ectopic nuclear TFEB localization in cardiomyocytes and leads to hypertrophic cardiomyopathy with features reminiscent of those observed in HAYOS patients carrying the ATAD3A p.R528W variant.^12, 66^ In addition, RagA/B-deficient mouse embryonic fibroblasts exhibit reduced lysosomal levels of v- ATPase V_1_ proteins,^66^ similar to the lysosomal proteomic defects observed in ATAD3A p.R528W-expressing cells. Consistent with this link, RagA-B promotes assembly of the active v- ATPase in *Drosophila* ovarian germline cells, ^70^ suggesting that reduced lysosomal RagA/B may contribute not only to aberrant TFEB activation but also to impaired lysosomal integrity and function in *ATAD3A* mutant cells.

The rescue of lysosomal expansion and developmental defects by *Mitf* knockdown indicates that aberrant TFEB/Mitf activation is a critical downstream consequence of pathogenic *ATAD3A* dysfunction. Under physiological conditions, transient TFEB/TFE3 activation promotes adaptive stress responses by expanding lysosomal capacity and enhancing degradation, autophagy, and mitophagy.^30^ In contrast, sustained TFEB/TFE3 activation can become maladaptive, disrupting lysosomal and organelle homeostasis and contributing to disease states, including tumorigenesis.^37^ This concept is supported by studies of tuberous sclerosis complex (TSC), in which loss-of-function mutations in *TSC1* or *TSC2* cause multisystem hamartomas and prominent neurological manifestations, accompanied by increased nuclear TFEB/TFE3 in mouse tissues and cultured human cells.^61^ Notably, kidney-specific *Tfeb* knockout rescues renal pathology in a mouse model of tuberous sclerosis complex.^62^ In addition, ectopic expression of *Tfeb* suppresses neuronal differentiation of neural progenitor cells in mice.^71^ Together, these findings support a model in which pathogenic *ATAD3A* variants drive disease by inducing chronic TFEB/Mitf activation that in turn perturbs lysosomal homeostasis during neurodevelopment.

We acknowledge several limitations in our study. First, we are unable to explain how ATAD3A ATP/ADP-bound states regulate coiled-coil topology and Rag GTPase biding. Because coiled-coil domain is also required for ATAD3A oligomerization,^13, 22, 72^ we cannot exclude the possibility that ATPase-dependent changes in oligomerization contribute to altered Rag GTPase interactions. In addition, the mechanisms by which impaired lysosomal localization of Rags and how chronic TFEB/Mitf activation drive neurodevelopmental phenotypes remain undefined. Future studies will define these downstream pathways and lysosomal dysfunction caused by ATAD3A mutations and test pharmaceutical interventions that may reverse ATAD3A p.R528W- mediated disruption of the mTOR-TFEB axis.

In summary, we report previously unrecognized effects of the pathogenic ATAD3A mutation on the RagC/D-TFEB axis in flies and human cells. ATAD3A interacts with RagC/D through its coiled-coil domain, and the p.R528W variant enhances this interaction while reducing lysosomal Rag GTPase localization. In human cells and *Drosophila*, ATAD3A p.R528W/dAtad3a p.R534W impairs mTORC1 substrate phosphorylation, promotes TFEB/Mitf activation, and drives lysosomal expansion. Deleting the coiled-coil domain abolishes these effects, while RagC-D expression or *Mitf* knockdown rescues neurodevelopmental and lysosomal phenotypes in vivo. We propose that pathogenic ATAD3A sequesters or mislocalizes Rag GTPases, leading to impaired mTORC1 signaling, chronic TFEB/Mitf activation, and disease-relevant lysosomal dysfunction. Our data provide compelling evidence highlighting cross-species genetic approaches can provide a foundation for unraveling potential pathogenic mechanisms.

## ACKNOWLEDGMENTS

W.H.Y. is supported by the National Institute of Neurological Disorders and Stroke (5R01 NS121298) of the National Institutes of Health (NIH). W.H.Y. is also supported by Presbyterian Health Foundation (PHF 4411-09-10-0) and Oklahoma Center for Adult Stem Cell Research (221009, 241006). M.B.M is supported by the Geroscience Training Grant (NIA T32AG052363). M.K. was supported by GM103447 and GM137786 of the NIH. S. L. is supported by the National Institute of Allergy and Infectious Diseases (R01-AI198430). We acknowledge the support from the OMRF Imaging Core Facility (P30-GM149376, National Institute of General Medical Sciences, NIH). We thank S. Plafker, I. Etchegaray, and J.S. Kang for their critical reading of the manuscript and helpful discussions. We acknowledge K.V. Parra, and Y. Park for technical assistance. Flybase, the Bloomington *Drosophila* Stock Center, and Vienna *Drosophila* Resource Center provided critical information and reagents for this study.

## AUTHOR CONTRIBUTIONS

M.B.M planned experimental design, conducted experiments, analyzed/visualized data, and wrote the manuscript. A.S. conducted experiments and analyzed/visualized data. M.K., A.J. and S.Y.J conducted mass spectrometry experiments. S. L. performed 3D structure modeling. W.H.Y conceived of the project, participated in experimental design, discussion of results and interpretation, and wrote the manuscript.

## COMPETING FINANCIAL INTERESTS

The authors declare no competing financial interests.

## Figure Legends

**Extended Figure 1. Expression of ATAD3A WT in human cells results in minimal effects on the proteome and transcriptome.**

(a) Volcano plot comparing proteomics analysis of SH-SY5Y cells expressing ATAD3A WT-V5 or Empty control. Blue represents less abundant (log_2_FC of <-0.263 and p-val <0.05) while orange represents more abundant (log_2_FC of >0.263 and p-val <0.05) proteins in ATAD3A WT-expressing cells. Results are from n=3 independent experiments, with 2 technical replicates run for each. (b) Volcano plot comparing RNA-Seq analysis of SH-SY5Y cells expressing ATAD3A WT-V5 or Empty control. Blue represents downregulated (log_2_FC of <-0.585 and p_adj_ <0.05) while orange represents upregulated (log_2_FC of >0.585 and p_adj_ <0.05) genes in ATAD3A WT- expressing cells. Results are from n=3 independent replicates.

**Extended Figure 2. Neuronal expression of dAtad3a p.R534W lacking coiled-coil domain did not cause developmental defects.**

Table showing lethality of flies expressing *ΔCC-dAtad3a^WT^*, *dAtad3a^R534W^*, and *ΔCC- dAtad3a^R534W^* in neurons.

**Extended Figure 3. ATAD3A lacking coiled-coil domain localizes to mitochondria and has minimal effects on lysosomal mTOR localization**

(a) Western blot for p18, p14, and TFEB in SH-SY5Y non-targeting control cells and *p18/LAMTOR1* knockout cells. (b) Confocal images of SH-SY5Y expressing ATAD3A WT, ATAD3A p.R528W, or ATAD3A ΔCC p.R528W. V5 (green) labels ATAD3A. COXIV (red) labels mitochondria. Scale bar, 20 μm. (c) Confocal images of SH-SY5Y p18 KO cells or control cells expressing Empty vector, ATAD3A WT, ATAD3A p.R528W, or ATAD3A ΔCC p.R528W stained for mTOR (green) and LAMP2 (red). Scale bar, 20 μm. Quantification of mTOR-positive LAMP2 puncta. Graphs represent mean +/- SEM from at least 15 images. p values were calculated using ordinary one-way ANOVA with Tukey’s multiple-comparisons test.. ns ≥ 0.05, \**P* < 0.05, and \*\*\*\**P* < 0.0001.

## METHODS

### *Drosophila* strains and maintenance

*UAS-dAtad3a-V5 lines including dAtad3a^WT^-V5* and *UAS-dAtad3a^p.R534W^-V5* and *dAtad3a-T2A- Gal4* line were generated as previously described.^26^ *elav-Gal4* (BL#458), *insc-Gal4* (BL#8751), UAS-witld-type RagC-D (BL# 83628), *Mitf* RNAi (BL#34835) and *UAS-GFP-LAMP* (BL# 42714) were obtained from the Bloomington *Drosophila* Stock Center. *asense-Gal4* lines and *UAS-Mitf- Myc* were generously provided from Tzumin Lee (Janelia Research Campus) and Francesca Pignoni (Upstate Medical University, NY), respectively. All flies were maintained at room temperature (21°C). Crosses were kept at 25°C.

### Cloning and transgenesis

*For cloning of pUASTattB-RagA-B,* cDNA (GH04846) obtained from the *Drosophila* Genomics Resource Center were cloned into pUASTattB vector using conventional restriction enzyme method, as previously described.^73^ pUASTattB constructs containing amino acid substitutions, including aAtad3a (p.G361D), aAtad3a (p.418Q), RagA-B (T16N), RagA-B (Q61L), RagC-D(S54N), or RagC-D(Q99L) were generated through site-directed mutagenesis, as previously described.^26^ The dAtad3a constructs were injected into y,w,ΦC31; VK37 and the Rag constructs were injected into y,w,ΦC31; attP2 embryos, and the transgenic flies were selected.

### Dissection and Immunostaining

Third instar larval brains were fixed in 4% formaldehyde for 30 min at room temperature and washed in PBS containing 0.3% Triton X-100. The primary antibodies were used at the following dilutions: Rat anti-Elav (7E8A10) (1:500) (DSHB Cat# Rat-Elav-7E8A10 anti-elav, RRID:AB_528218), Ms anti-Prospero (MR1A) (1:500) (DSHB Cat# Prospero, RRID:AB_528440), Rb anti-Myc (1:50) (Santa Cruz Biotechnology Cat# sc-789-G, RRID:AB_631275), Ms anti-Lamin C (ADL84.12) (1:100) (DSHB Cat# adl84.12, RRID:AB_528338), Rat anti-Miranda (1:500) (Abcam Cat# ab197788, RRID:AB_2936368), Rb anti-GFP (1:1000) (Molecular Probes Cat# A-11122, RRID:AB_221569), Rt anti-Miranda (1:500) (Abcam Cat# ab197788, RRID:AB_2936368), Ms Elav 9F8A9 (1:500) (DSHB Cat# Elav-9F8A9, RRID:AB_528217). Secondary antibodies were used at the following dilutions: Goat anti-Ms Cy5 (1:500) (Thermo Fisher Scientific Cat# A10524, RRID:AB_10562712), Donkey anti-Rat 488 (1:500) (Molecular Probes Cat# A-21208, RRID:AB_2535794), Goat anti-Ms Alexa Fluor 568 (Thermo Fisher Scientific Cat# A-11004, RRID:AB_2534072), Donkey anti-Rb Alexa Fluor 488 (Thermo Fisher Scientific Cat# A-21206, RRID:AB_2535792), Goat anti-Rt 568 (1:500) (Thermo Fisher Scientific Cat# A-11077, RRID:AB_2534121), and Donkey anti-Ms 647 (1:250) Jackson Lab (Jackson ImmunoResearch Labs Cat# 715-605-150, RRID:AB_2340862). Samples were mounted in Vectashield (Vector Labs, Burlingame, CA). Imaging was performed using LSM880 confocal microscope (Zeiss). Images were processed with Zeiss LSM Image Browser and Adobe Photoshop. Intensity of GFP-LAMP and Mitf-Myc was quantified using ImageJ.

### Modeling of ATAD3A AAA+ structure

The AlphaFold-predicted model (AF-Q9NVI7-F1) was used as the starting structure.^41^ To add the missing ATP molecule, the crystal structure of the C-terminal domain of ClpB (PDB ID: 4FCV, chain A) was superimposed onto the model. Due to domain motion between the large αβ ATP binding domain and the small helical domain, overall fitting was not optimal. Therefore, only the large αβ domain was used for the superposition, and AMP-PNP was subsequently modeled into the AlphaFold-predicted ATAD3A structure. To model the Tryptophan side chain introduced by the p.Arg528Trp mutation, the program Coot was used,^42^ and a rotamer closely matching the original arginine side-chain orientation was selected.

### Cell Culture and Cell Lines

All cells were incubated at 37°C and 5% CO_2_ and confirmed Mycoplasma negative by PCR (Sigma MP0025). SH-SY5Y cells were grown in DMEM (Gibco #10569010) supplemented with 15% heat-inactivated FBS (SA F0926), 1x NEAA (Gibco # 11140050), and 1x Pen/Strep (Gibco #15070063). HEK293T cells were grown in DMEM (Gibco #10569010) supplemented with 10% heat-inactivated FBS (SA F0926), and 1x Pen/Strep (Gibco # 15070063).

### Plasmids and Cloning

Plasmids pRK5-HA GST RAP2A (Addgene plasmid #14952; http://n2t.net/addgene:14952; RRID:Addgene_14952), pRK5-HA GST RHEB (Addgene plasmid #14951; http://n2t.net/addgene:14951; RRID:Addgene_14951), pRK5-HA GST RagA wt (Addgene plasmid # 19298; http://n2t.net/addgene:19298; RRID:Addgene_19298), pRK5-HA GST RagB wt (Addgene plasmid #19301; http://n2t.net/addgene:19301; RRID:Addgene_19301), pRK5-HA GST RagC wt (Addgene plasmid #19304; http://n2t.net/addgene:19304; RRID:Addgene_19304), pRK5-HA GST RagD wt (Addgene plasmid #19307; http://n2t.net/addgene:19307; RRID:Addgene_19307), FLAG pLJM1 RagD (Addgene plasmid #19316; http://n2t.net/addgene:19316; RRID:Addgene_19316), pLJC5 TMEM192 2xFLAG (Addgene plasmid #102929; http://n2t.net/addgene:102929; RRID:Addgene_102929), and pLJC5 TMEM192 3xHA (Addgene plasmid #102930; http://n2t.net/addgene:102930; RRID:Addgene_102930) were gifts from David Sabatini. pLJM1-Empty was a gift from Joshua Mendell (Addgene plasmid #91980; http://n2t.net/addgene:91980; RRID:Addgene_91980). FLAG pLJM1 RagC WT was generated by PCR of pRK5 Flag-RagC (gift from David Sabatini & Kuang Shen [Addgene plasmid # 99723; http://n2t.net/addgene:99723; RRID:Addgene 99723]) followed by restriction enzyme digestion and ligation in the previously mentioned FLAG pLJM1 RagD vector, replacing the RagD coding sequence for RagC.

pcDNA3.1 ATDA3A constructs containing amino acid substitutions, including ATAD3A p.G355D, p.K358A, p.R528W, and p.E412Q were generated through site-directed mutagenesis, as previously described.^12^ pcDNA3.1 ATAD3A and ATAD3A p.R528W were PCR amplified and cloned into the LentiV_Blast vector, which was a gift from Christopher Vakoc (Addgene plasmid # 111887; http://n2t.net/addgene:111887; RRID:Addgene_111887), using restriction enzyme cloning.

Generation of coiled-coil deletions in ATAD3A or dAtad3a were performed by restriction digest followed by Gibson assembly of a synthesized gBlock (Integrated DNA Technologies) using the Gibson Assembly Master Mix (New England BioLabs E2611S) into the parental vector following manufacturer instructions.

LentiCRISPRv2 plasmids were generated as previously described using a scramble control (5’- GCACTACCAGAGCTAACTCA-3’) or LAMTOR1-targeting sequence (5’- GTGTCTGCTGCAGACTCACA-3’) obtained from the Brunello library.^74^

### Lentivirus production and transduction

HEK293Ts were plated on poly-L-lysine (Sigma P4707) coated 6w plates at 400,000 c/w. The following day, HEK293Ts were co-transfected with Lentiviral vectors and packaging plasmids pVSV-G and pCMVR8.74 in 1mL of 3% FBS containing DMEM. After 6h, media was changed to 1.5mL DMEM supplemented with 3% FBS and 1x NEAA. 48h post-transfection, supernatants were harvested and stored at 4°C overnight. 1.5mL of fresh DMEM supplemented with 3% FBS and 1x NEAA was added to lentivirus-producing cells. 72h post-transfection, lentivirus- containing supernatants were added to the 48h supernatant and HEPES and Polybrene were added to a final concentration of 20mM and 4µg/mL, respectively. Supernatants were then filtered through 0.45µm filters, aliquoted, and frozen at −80°C.

Before transduction, SH-SY5Y cells were plated at a density of 300,000 c/w. The next day, media was changed to 1mL of DMEM containing 3% FBS, 20mM HEPES, and 4µg/mL Polybrene and Lentivirus was added to each well. 6h later, 1mL of DMEM supplemented with 15% FBS and 1x NEAA was added. 48h post-transduction, cells were harvested with Trypsin and passaged into selection-containing media (Blasticidin: 12µg/mL and/or Puromycin: 1µg/mL).

### Immunohistochemistry of human cells

Cells were plated in a 6w plate containing Poly-L-lysine (P4707) coated coverglass. On day 4 after plating, cells were washed 2x with PBS and then fixed in 4% PFA (Electron Microscopy Sciences 15710) in PBS for 10min. After being washed with PBS and permeabilized with PBS-T 1x PBS with 0.1% TritonX-100 (PBS-T), samples were blocked with 5% Norman Goat Serum (Jackson Labs #005-000-121) in PBS-T for at least 1h. After blocking, cover glass was removed from plates and incubated in primary antibody (Ms anti-LAMP2 [Santa Cruz Biotechnology Cat# sc-18822, RRID:AB_626858] 1:50, Rb anti-RagC [Cell Signaling Technology Cat# 9480, RRID:AB_10614716] 1:50, and/or Rb anti-mTOR [Cell Signaling Technology Cat# 2983, RRID:AB_2105622] 1:100) with cover glass face down on parafilm containing primary antibody diluted in 5% Norman Goat Serum in PBS-T overnight at 4°C. The next day, cover glass with placed back into the 6w plate and washed 4x with PBS-T for 15min each. For mitochondrial localization experiments, cover glass was incubated in primary antibody within 6w plates at 1:500 Rb anti-COXIV (Cell Signaling Technology Cat# 4850, RRID:AB_2085424) and 1:500 Ms anti-V5 (Thermo Fisher Scientific Cat# R960-25, RRID:AB_2556564) rocking overnight followed by 4x washes with PBS-T. After washing, samples were incubated with secondary antibody Donkey anti-Rb Alexa Fluor 488 (Thermo Fisher Scientific Cat# A-21206, RRID:AB_2535792), Goat anti-Rb Alexa Fluor 568 (Thermo Fisher Scientific Cat# A-11011, RRID:AB_143157), Donkey anti-Ms Alexa Fluor 488 (Thermo Fisher Scientific Cat# A-21202, RRID:AB_141607), or Goat anti-Ms Alexa Fluor 568 (Thermo Fisher Scientific Cat# A-11004, RRID:AB_2534072) diluted 1:500 in PBS-T for 1h. After rinsing with PBS-T, samples were incubated with DAPI (1µg/mL) for 20-30 minutes. Following 3x washes with PBS-T, samples were washed 1x with PBS and mounted on microscopy glass in Vectashield mounting media and sealed with clear nail polish.

Images were taken with a Zeiss LSM880 microscope and analyzed with Imaris software using the surface function. Co-localization between LAMP2 and RagC or mTOR was determined by calculating the number of LAMP2 surfaces with less than 0.2µm distance to a RagC or mTOR surface.

### Western blotting, Cell Fractionation, and Proteomics

Cells harvested with RIPA Buffer by scraping cells from plates kept on ice and transferred to tubes. Tubes containing lysate were then vortexed for 10s every 5 min for a total of 30min. Lysates were then spun at 4°C for 15 min at 20,817xg. Supernatant was aliquoted and stored at −80 C until SDS-PAGE.

For SDS-PAGE, loading of lysates was normalized by Bradford assay and run on precast BioRad gradient gels. After SDS-PAGE, protein was transferred using the BioRad TransTurbo transfer system onto nitrocellulose membranes. Membranes were then blocked for at least 45 minutes with 5% Milk in TBS-T, briefly washed with TBS-T, and incubated in primary antibody diluted in 5% BSA in TBS-T overnight. The following day, membranes were washed with TBS-T 3x for 10min. Secondary antibody was then applied at 1:5000 (Goat anti-Rb HRP (Thermo Fisher Scientific Cat# G-21234, RRID:AB_2536530) or Goat anti-Ms HRP (Thermo Fisher Scientific Cat# A28177, RRID:AB_2536163) in 5% milk in TBS-T for at least 1h, followed by washes with TBS-T 3x for 10min. Bands were then visualized by application of Clarity ECL substrate (BioRad #170-5060) for 1-2 minutes and imaged using the BioRad ChemiDoc MP imaging system.

The following antibodies were utilized for Western blotting: Rb anti-TFEB (Cell Signaling Technology Cat# 4240, RRID:AB_11220225), Rb anti-S6 ribosomal protein (Cell Signaling Technology Cat# 2217, RRID:AB_331355), Rb anti-phospho S6 ribosomal protein (Ser235/236) (Cell Signaling Technology Cat# 2211, RRID:AB_331679), Rb anti-4E-BP1 (Cell Signaling Technology Cat# 9644, RRID:AB_2097841), Rb anti-phospho 4E-BP1 (Thr37 / Thr46) (Cell Signaling Technology Cat# 2855, RRID:AB_560835), Rb anti-p70 S6 Kinase (Cell Signaling Technology Cat# 9202, RRID:AB_331676), Rb anti-phospho-p70 S6 Kinase (Thr389) (Cell Signaling Technology Cat# 97596, RRID:AB_2800283), Rb anti-ATAD3A (Novus Cat# H00055210-D01P, RRID:AB_11027573), Rb anti-RagA (Cell Signaling Technology Cat# 4357, RRID:AB_10545136), Rb anti-RagB (Cell Signaling Technology Cat# 8150, RRID:AB_11178806), Rb anti-RagC (Cell Signaling Technology Cat# 9480, RRID:AB_10614716), Rb anti-RagD (Bethyl Cat# A304-301A, RRID:AB_2620497), Rb anti- LAMTOR1/p18 (Cell Signaling Technology Cat# 8975, RRID:AB_10860252), Rb anti- LAMTOR2/p14 (Cell Signaling Technology Cat# 8145, RRID:AB_10971636), Rb anti-mTOR (Cell Signaling Technology Cat# 2983, RRID:AB_2105622), Rb anti-TFE3 (Proteintech Cat# 14480-1-AP, RRID:AB_2199587), Rb anti-Lamin A/C (Cell Signaling Technology Cat# 2032, RRID:AB_2136278), Rb anti-ATP6V1A (Abcam Cat# ab199326, RRID:AB_2802119), Rb anti- ATP6V0D1 (Abcam Cat# ab202899),Ms anti-GAPDH (Proteintech Cat# 60004-1-Ig, RRID:AB_2107436), Ms anti-LAMP2 (Santa Cruz Biotechnology Cat# sc-18822, RRID:AB_626858), Ms anti-Actin (MP Bio Cat# 0869100-CF, RRID:AB_2920628), and Ms anti- V5 (Thermo Fisher Scientific Cat# R960-25, RRID:AB_2556564).

Cell fractionation experiments were performed using the cell fractionation kit (Cell Signaling Technology #9038) following manufacturer instructions modifying the sonication steps with use of a Biorupter plus (Diagenode) on the high setting.

Proteomics analysis was performed on samples harvested in RIPA buffer as described above. Protein lysate was subjected to SDS-PAGE followed by trypsin digestion at room temperature overnight. Peptides were extracted with 50% acetonitrile, dried by Speedvac, and reconstituted in 200µL 1% acetic acid. Data independent acquisition was then performed using an Orbitrap Exploris 480 Mass Spectrometer. Data were analyzed using DIA-NN^75^ and normalized to a horse serum albumin internal standard.

Gene set enrichment analysis was performed using the Molecular Signatures Database (MSigDB) Hallmark gene set collection^45^ using g:Profiler.^76^

### Lysosome Immunoprecipitation (Lyso-IP) and Mass Spectrometry

Lyso-IP was slightly modified from previous studies^53^. All centrifuge steps were done at 4°C. Briefly, cells were washed with ice-cold PBS, scraped into KPBS on ice, normalized by counting, and spun at 1000xg for 2 min. Cells were resuspended in 950uL of KPBS and homogenized with a 2mL Dounce homogenizer with 10 loose, followed by 15 tight strokes on ice. Lysate was then transferred to a tube and spun at 1000xg for 2 min. The supernatant was transferred to prewashed anti-HA magnetic beads (ThermoFisher 88837) and rotated for 30 min at 4°C. Tubes were then placed into a magnetic rack for 30s, supernatant was removed, and beads were resuspended with 1mL of cold KPBS all inside of a cold room at 4°C. This was repeated 1x with KPBS, and then 1x with KPBS containing 150mM NaCl. Following resuspension, beads were transferred to a fresh tube and washed a final time in KPBS at 4°C. Proteins were then eluted from magnetic beads with 0.5% NP-40 for 30 min rotating at 4°C. Following a 30 second incubation on the magnetic rack, the elution supernatant was moved to a new tube. Protein eluates were then flash frozen in liquid nitrogen and stored at −80°C until downstream analyses.

For proteomics, protein eluates were run by SDS-PAGE and trypsin digested overnight at room temperature. Peptides were extracted from the gel with 50% acetonitrile, dried by Speedvac, and reconstituted in 200µL 1% acetic acid. Data independent acquisition was then performed using an Orbitrap Exploris 480 Mass Spectrometer. Data were analyzed using DIA-NN^75^, and each sample was normalized to its own total ion current.

### Immunoprecipitation/Mass Spectrometry

The protein lysate was incubated with 2µg of V5 antibody for 1h followed by ultra-centrifugation at 100,000g, 20min, 4°C and 1hour incubation with Sepharose-CL4B Protein A beads (GE Healthcare). The beads were washed three times with lysis buffer (150mM NaCl, 50mM Tris-Cl pH 8.0, 1mM EDTA, 0.5% NP-40). The protein complex was eluted using 1X NuPAGE LDS sample buffer (Invitrogen NP0007), resolved on 10% Bis-Tris NuPAGE gel (Invitrogen NP0315BOX), and visualized with Coomassie Brilliant blue-stain. The whole lane was in-gel digested with trypsin enzyme as described previously^77^ to obtain 2 peptide pools. The peptide pools were concentrated in a speed vac and dissolved in 5% methanol containing 0.1% formic acid loading solution. The LC-MS/MS analysis was carried out on a nano-LC 1000 system (Thermo Fisher Scientific, San Jose, CA) coupled to Orbitrap Q-Exactive Plus (Thermo Fisher Scientific, San Jose, CA). The peptides were eluted using a 55min gradient of 4-26% acetonitrile/0.1% formic acid at a flow rate of 800nl/min followed by a 5min wash. The mass spectrometer was operated in the data-dependent acquisition mode with top35 dependent scans. MS1 was acquired in Orbitrap (140,000 resolution, 375-1300m/z) followed by MS2 in Orbitrap (17,500 resolution, 30ms injection time). The MS raw data was searched in Proteome Discoverer software (v1.4, Thermo Scientific, San Jose, CA) with Mascot algorithm (v2.4, Matrix Science,)^78^ against the *Drosophila melanogaster* NCBI refseq database (updated 2019_01_14). The precursor and fragment mass tolerance were set to 20ppm and 0.02Da respectively. Maximum miscut of 2 with trypsin enzyme, dynamic modification of oxidation (M), protein N-term acetylation, and destreak (C) was allowed. The peptides identified from mascot result file were validated with 5% false discover rate (FDR) in Percolator.^79^ The PSMs result file was further processed through ‘gpGrouper’ algorithm.^80^ The protein-level inference and quantification was conducted with label-free iBAQ approach using ‘gpGrouper’ algorithm. The log-transformed iBAQ values were used for differential analysis. Differential expression analysis was performed using the limma moderated t-test, and log2 fold changes were calculated using the limma package in R.

### RNA Isolation and RNA-Seq

To harvest RNA, cells were treated with TRIzol reagent (Invitrogen #15596026) according to manufacturer instructions and stored at −80°C until RNA isolation. RNA isolated with the RNA Clean & Concentrator −25 kit (Zymo R1017) according to manufacturer instructions. mRNA enrichment, library preparation, and RNA-Seq was performed by Novogene. Reads were mapped to the human genome and differentially expressed gene analysis was performed using DeSEQ2. Gene set enrichment analysis was performed using the Molecular Signatures Database (MSigDB) Hallmark gene set collection^45^ using g:Profiler.^76^

### Statistical Analysis

Statistical analysis was performed using Microsoft Excel or GraphPad Prism 11.

## Notes

### Competing Interest Statement

The authors have declared no competing interest.

## REFERENCES

1. DiMauro, S. & Schon, E.A. Mitochondrial disorders in the nervous system. Annu Rev Neurosci 31, 91–123 (2008).

2. Fraldi, A., Klein, A.D., Medina, D.L. & Settembre, C. Brain Disorders Due to Lysosomal Dysfunction. Annu Rev Neurosci 39, 277–295 (2016).

3. Prashar, A. & Puertollano, R. Neighbors who talk: Mitochondria-lysosome crosstalk in homeostasis. Curr Opin Cell Biol 100, 102627 (2026).

4. Demers-Lamarche, J. et al. Loss of Mitochondrial Function Impairs Lysosomes. J Biol Chem 291, 10263–10276 (2016).

5. Kim, S., Wong, Y.C., Gao, F. & Krainc, D. Dysregulation of mitochondria- lysosome contacts by GBA1 dysfunction in dopaminergic neuronal models of Parkinson’s disease. Nat Commun 12, 1807 (2021).

6. He, J. et al. The AAA+ protein ATAD3 has displacement loop binding properties and is involved in mitochondrial nucleoid organization. J Cell Biol 176, 141–146 (2007).

7. Gilquin, B. et al. The AAA+ ATPase ATAD3A controls mitochondrial dynamics at the interface of the inner and outer membranes. Mol Cell Biol 30, 1984–1996 (2010).

8. Issop, L. et al. Mitochondria-associated membrane formation in hormone- stimulated Leydig cell steroidogenesis: role of ATAD3. Endocrinology 156, 334–345 (2015).

9. Desai, R. et al. ATAD3 gene cluster deletions cause cerebellar dysfunction associated with altered mitochondrial DNA and cholesterol metabolism. Brain 140, 1595–1610 (2017).

10. Munoz-Oreja, M. et al. Elevated cholesterol in ATAD3 mutants is a compensatory mechanism that leads to membrane cholesterol aggregation. Brain 147, 1899–1913 (2024).

11. Waters, E.R., Bezanilla, M. & Vierling, E. ATAD3 Proteins: Unique Mitochondrial Proteins Essential for Life in Diverse Eukaryotic Lineages. Plant Cell Physiol 65, 493–502 (2024).

12. Harel, T. et al. Recurrent De Novo and Biallelic Variation of ATAD3A, Encoding a Mitochondrial Membrane Protein, Results in Distinct Neurological Syndromes. Am J Hum Genet 99, 831–845 (2016).

13. Rigoni, G. et al. MARIGOLD and MitoCIAO, two searchable compendia to visualize and functionalize protein complexes during mitochondrial remodeling. Cell Metab 37, 1024–1038 e1028 (2025).

14. Hu, C. et al. OPA1 and MICOS Regulate mitochondrial crista dynamics and formation. Cell Death Dis 11, 940 (2020).

15. Arguello, T. et al. ATAD3A has a scaffolding role regulating mitochondria inner membrane structure and protein assembly. Cell Rep 37, 110139 (2021).

16. Peralta, S. et al. ATAD3 controls mitochondrial cristae structure in mouse muscle, influencing mtDNA replication and cholesterol levels. J Cell Sci 131 (2018).

17. Ishihara, T., Ban-Ishihara, R., Ota, A. & Ishihara, N. Mitochondrial nucleoid trafficking regulated by the inner-membrane AAA-ATPase ATAD3A modulates respiratory complex formation. Proc Natl Acad Sci U S A 119, e2210730119 (2022).

18. Jin, G. et al. Atad3a suppresses Pink1-dependent mitophagy to maintain homeostasis of hematopoietic progenitor cells. Nat Immunol 19, 29–40 (2018).

19. Frazier, A.E. et al. Fatal perinatal mitochondrial cardiac failure caused by recurrent de novo duplications in the ATAD3 locus. Med 2, 49–73 (2021).

20. Bae, T., et al. Allele-specific correction of ATAD3A pathogenic variants via template-free CRISPR-Cas9 editing and gene conversion. bioRxiv (2025).

21. Baudier, J. ATAD3 proteins: brokers of a mitochondria-endoplasmic reticulum connection in mammalian cells. Biol Rev Camb Philos Soc 93, 827–844 (2018).

22. Zhao, Y. et al. ATAD3A oligomerization causes neurodegeneration by coupling mitochondrial fragmentation and bioenergetics defects. Nat Commun 10, 1371 (2019).

23. Brar, K.K. et al. PERK-ATAD3A interaction provides a subcellular safe haven for protein synthesis during ER stress. Science 385, eadp7114 (2024).

24. Goller, T., Seibold, U.K., Kremmer, E., Voos, W. & Kolanus, W. Atad3 function is essential for early post-implantation development in the mouse. PLoS One 8, e54799 (2013).

25. Ezer, S. et al. Transcriptome analysis of atad3-null zebrafish embryos elucidates possible disease mechanisms. Orphanet J Rare Dis 20, 181 (2025).

26. Yap, Z.Y. et al. Functional interpretation of ATAD3A variants in neuro- mitochondrial phenotypes. Genome Med 13, 55 (2021).

27. Hoffmann, M. et al. C. elegans ATAD-3 is essential for mitochondrial activity and development. PLoS One 4, e7644 (2009).

28. Brugel, M., Kiesel, A.S., Haack, T.B. & Peralta, S. Mutations in mitochondrial ATAD3 gene and disease, lessons from in vivo models. Front Neurosci 18, 1496142 (2024).

29. Cooper, H.M. et al. ATPase-deficient mitochondrial inner membrane protein ATAD3A disturbs mitochondrial dynamics in dominant hereditary spastic paraplegia. Hum Mol Genet 26, 1432–1443 (2017).

30. Raben, N. & Puertollano, R. TFEB and TFE3: Linking Lysosomes to Cellular Adaptation to Stress. Annu Rev Cell Dev Biol 32, 255–278 (2016).

31. Sardiello, M. et al. A gene network regulating lysosomal biogenesis and function. Science 325, 473–477 (2009).

32. Roczniak-Ferguson, A. et al. The transcription factor TFEB links mTORC1 signaling to transcriptional control of lysosome homeostasis. Sci Signal 5, ra42 (2012).

33. Martina, J.A. et al. The nutrient-responsive transcription factor TFE3 promotes autophagy, lysosomal biogenesis, and clearance of cellular debris. Sci Signal 7, ra9 (2014).

34. Medina, D.L. et al. Lysosomal calcium signalling regulates autophagy through calcineurin and TFEB. Nat Cell Biol 17, 288–299 (2015).

35. Palmieri, M. et al. Characterization of the CLEAR network reveals an integrated control of cellular clearance pathways. Hum Mol Genet 20, 3852–3866 (2011).

36. Martina, J.A. & Puertollano, R. Rag GTPases mediate amino acid-dependent recruitment of TFEB and MITF to lysosomes. J Cell Biol 200, 475–491 (2013).

37. Napolitano, G. et al. A substrate-specific mTORC1 pathway underlies Birt-Hogg- Dube syndrome. Nature 585, 597–602 (2020).

38. Sancak, Y. et al. Ragulator-Rag complex targets mTORC1 to the lysosomal surface and is necessary for its activation by amino acids. Cell 141, 290–303 (2010).

39. Villegas, F. et al. Lysosomal Signaling Licenses Embryonic Stem Cell Differentiation via Inactivation of Tfe3. Cell Stem Cell 24, 257–270 e258 (2019).

40. Chen, D. et al. Mitochondrial ATAD3A regulates milk biosynthesis and proliferation of mammary epithelial cells from dairy cow via the mTOR pathway. Cell Biol Int 42, 533–542 (2018).

41. Jumper, J. et al. Highly accurate protein structure prediction with AlphaFold. Nature 596, 583–589 (2021).

42. Emsley, P., Lohkamp, B., Scott, W.G. & Cowtan, K. Features and development of Coot. Acta Crystallogr D Biol Crystallogr 66, 486–501 (2010).

43. Hanson, P.I. & Whiteheart, S.W. AAA+ proteins: have engine, will work. Nat Rev Mol Cell Biol 6, 519–529 (2005).

44. Dai, Y. et al. Cryo-EM structure of the AAA+ SPATA5 complex and its role in human cytoplasmic pre-60S maturation. Nat Commun 16, 3806 (2025).

45. Liberzon, A. et al. The Molecular Signatures Database (MSigDB) hallmark gene set collection. Cell Syst 1, 417–425 (2015).

46. Lepelley, A. et al. Enhanced cGAS-STING-dependent interferon signaling associated with mutations in ATAD3A. J Exp Med 218 (2021).

47. Wu, H. et al. Mitochondrial dysfunction promotes the transition of precursor to terminally exhausted T cells through HIF-1alpha-mediated glycolytic reprogramming. Nat Commun 14, 6858 (2023).

48. Di Malta, C. et al. Transcriptional activation of RagD GTPase controls mTORC1 and promotes cancer growth. Science 356, 1188–1192 (2017).

49. Carey, K.L. et al. TFEB Transcriptional Responses Reveal Negative Feedback by BHLHE40 and BHLHE41. Cell Rep 33, 108371 (2020).

50. Dubouloz, F., Deloche, O., Wanke, V., Cameroni, E. & De Virgilio, C. The TOR and EGO protein complexes orchestrate microautophagy in yeast. Mol Cell 19, 15–26 (2005).

51. Kim, E., Goraksha-Hicks, P., Li, L., Neufeld, T.P. & Guan, K.L. Regulation of TORC1 by Rag GTPases in nutrient response. Nat Cell Biol 10, 935–945 (2008).

52. Sancak, Y. et al. The Rag GTPases bind raptor and mediate amino acid signaling to mTORC1. Science 320, 1496–1501 (2008).

53. Abu-Remaileh, M. et al. Lysosomal metabolomics reveals V-ATPase- and mTOR- dependent regulation of amino acid efflux from lysosomes. Science 358, 807–813 (2017).

54. Li, F. & Zhang, H. Lysosomal Acid Lipase in Lipid Metabolism and Beyond. Arterioscler Thromb Vasc Biol 39, 850–856 (2019).

55. Zhang, H. Lysosomal acid lipase and lipid metabolism: new mechanisms, new questions, and new therapies. Curr Opin Lipidol 29, 218–223 (2018).

56. Yang, G. et al. RagC phosphorylation autoregulates mTOR complex 1. EMBO J 38 (2019).

57. Binda, M. et al. The Vam6 GEF controls TORC1 by activating the EGO complex. Mol Cell 35, 563–573 (2009).

58. Kim, W., Jang, Y.G., Yang, J. & Chung, J. Spatial Activation of TORC1 Is Regulated by Hedgehog and E2F1 Signaling in the Drosophila Eye. Dev Cell 42, 363–375 e364 (2017).

59. Romero-Pozuelo, J., Demetriades, C., Schroeder, P. & Teleman, A.A. CycD/Cdk4 and Discontinuities in Dpp Signaling Activate TORC1 in the Drosophila Wing Disc. Dev Cell 42, 376–387 e375 (2017).

60. Bar-Peled, L., Schweitzer, L.D., Zoncu, R. & Sabatini, D.M. Ragulator is a GEF for the rag GTPases that signal amino acid levels to mTORC1. Cell 150, 1196–1208 (2012).

61. Alesi, N. et al. TSC2 regulates lysosome biogenesis via a non-canonical RAGC and TFEB-dependent mechanism. Nat Commun 12, 4245 (2021).

62. Alesi, N. et al. TFEB drives mTORC1 hyperactivation and kidney disease in Tuberous Sclerosis Complex. Nat Commun 15, 406 (2024).

63. Bouche, V. et al. Drosophila Mitf regulates the V-ATPase and the lysosomal- autophagic pathway. Autophagy 12, 484–498 (2016).

64. Baixauli, F. et al. Mitochondrial Respiration Controls Lysosomal Function during Inflammatory T Cell Responses. Cell Metab 22, 485–498 (2015).

65. Cui, Z. et al. Structure of the lysosomal mTORC1-TFEB-Rag-Ragulator megacomplex. Nature 614, 572–579 (2023).

66. Kim, Y.C. et al. Rag GTPases are cardioprotective by regulating lysosomal function. Nat Commun 5, 4241 (2014).

67. Meireles, A.M. et al. The Lysosomal Transcription Factor TFEB Represses Myelination Downstream of the Rag-Ragulator Complex. Dev Cell 47, 319–330 e315 (2018).

68. Jewell, J.L. et al. Metabolism. Differential regulation of mTORC1 by leucine and glutamine. Science 347, 194–198 (2015).

69. Figlia, G. et al. Brain-enriched RagB isoforms regulate the dynamics of mTORC1 activity through GATOR1 inhibition. Nat Cell Biol 24, 1407–1421 (2022).

70. Zhou, Y. et al. Rag GTPases control lysosomal acidification by regulating v- ATPase assembly in Drosophila. J Biol Chem 301, 110400 (2025).

71. Yuizumi, N. et al. Maintenance of neural stem-progenitor cells by the lysosomal biosynthesis regulators TFEB and TFE3 in the embryonic mouse telencephalon. Stem Cells 39, 929–944 (2021).

72. Zhao, Y. et al. ATAD3A oligomerization promotes neuropathology and cognitive deficits in Alzheimer’s disease models. Nat Commun 13, 1121 (2022).

73. Yoon, W.H. et al. Loss of Nardilysin, a Mitochondrial Co-chaperone for alpha- Ketoglutarate Dehydrogenase, Promotes mTORC1 Activation and Neurodegeneration. Neuron 93, 115–131 (2017).

74. Doench, J.G. et al. Optimized sgRNA design to maximize activity and minimize off-target effects of CRISPR-Cas9. Nat Biotechnol 34, 184–191 (2016).

75. Demichev, V., Messner, C.B., Vernardis, S.I., Lilley, K.S. & Ralser, M. DIA-NN: neural networks and interference correction enable deep proteome coverage in high throughput. Nat Methods 17, 41–44 (2020).

76. Kolberg, L. et al. g:Profiler-interoperable web service for functional enrichment analysis and gene identifier mapping (2023 update). Nucleic Acids Res 51, W207–W212 (2023).

77. Chen, Y. et al. A Cross-Linking-Aided Immunoprecipitation/Mass Spectrometry Workflow Reveals Extensive Intracellular Trafficking in Time-Resolved, Signal- Dependent Epidermal Growth Factor Receptor Proteome. J Proteome Res 18, 3715–3730 (2019).

78. Perkins, D.N., Pappin, D.J., Creasy, D.M. & Cottrell, J.S. Probability-based protein identification by searching sequence databases using mass spectrometry data. Electrophoresis 20, 3551–3567 (1999).

79. Kall, L., Canterbury, J.D., Weston, J., Noble, W.S. & MacCoss, M.J. Semi- supervised learning for peptide identification from shotgun proteomics datasets. Nat Methods 4, 923–925 (2007).

80. Saltzman, A.B. et al. gpGrouper: A Peptide Grouping Algorithm for Gene-Centric Inference and Quantitation of Bottom-Up Proteomics Data. Mol Cell Proteomics 17, 2270–2283 (2018).

