## Supplementary figures and images for "Pathogenic mutations in ATAD3A cause dysregulation of RagC/D-TFEB axis and disrupt lysosomal homeostasis"

### Ext-Figure1_ATAD3A_BioRxiv(2026-09-23).tif

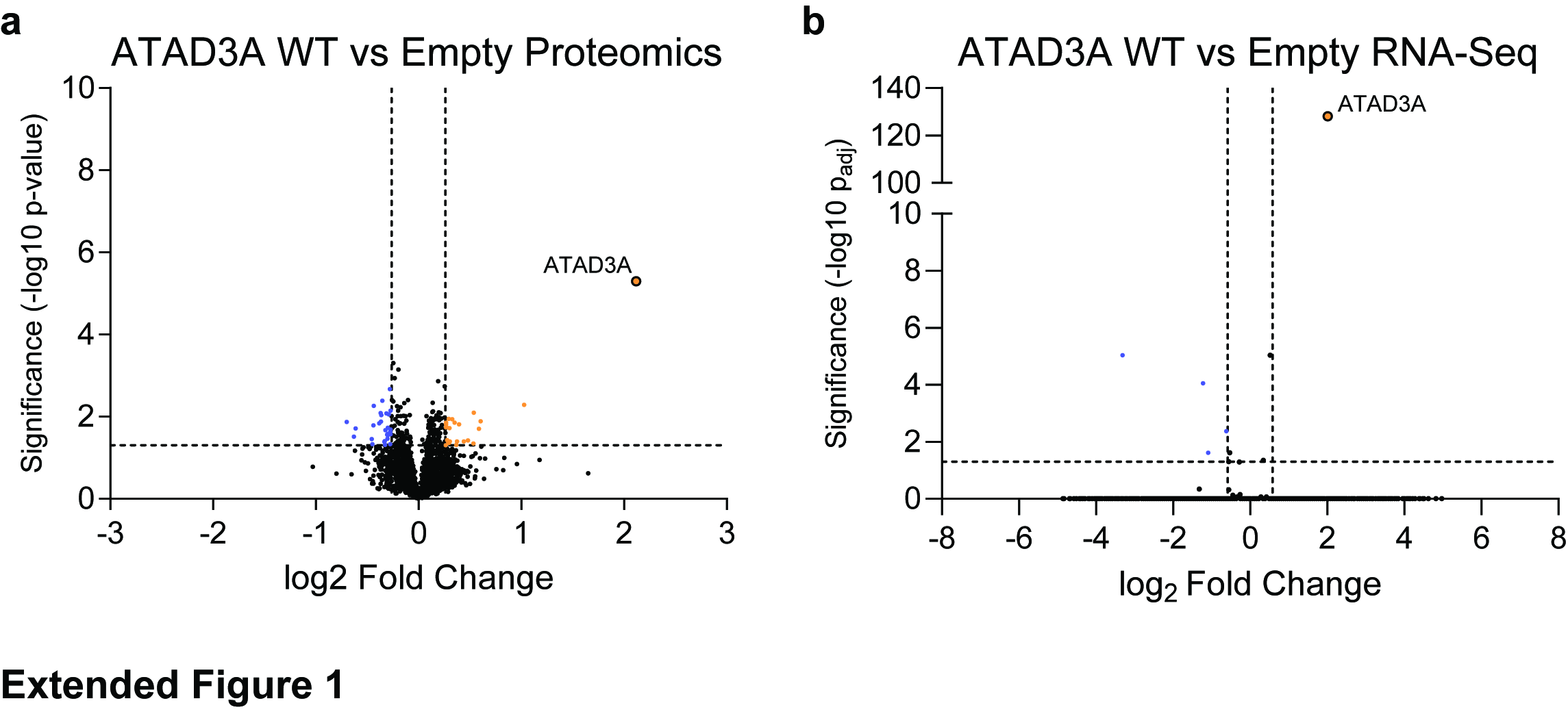

### Ext-Figure 3_ATAD3A_BioRxiv(2026-09-23).tif

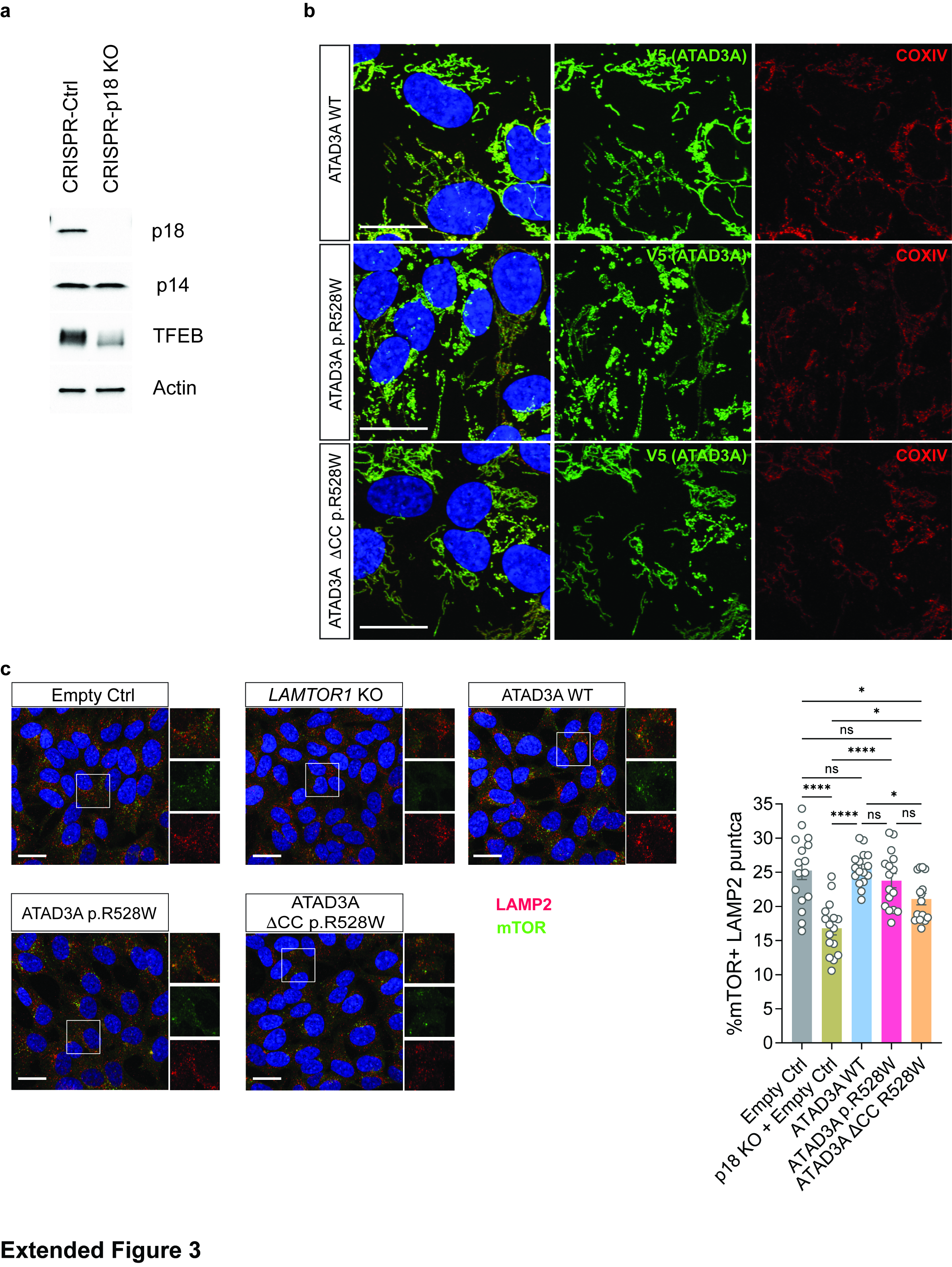

### Extended Figure 2

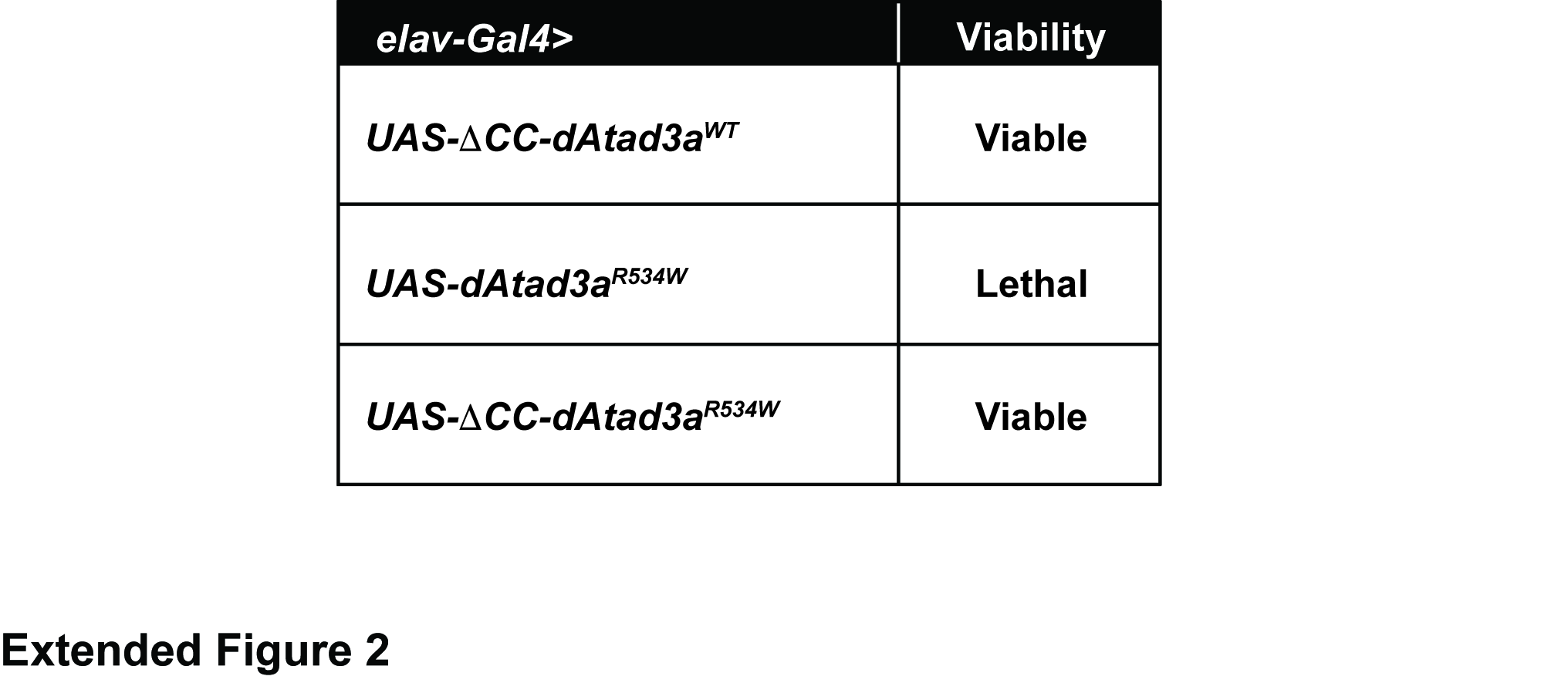
